# Clonal overlap and convergent clustering of T-cell receptor signatures in Crohn’s disease in monozygotic twins

**DOI:** 10.1101/2025.10.31.685913

**Authors:** Eelco C. Brand, Romi Vandoren, Lisanne Lutter, Vincent M.L. Van Deuren, Nila H. Servaas, Aridaman Pandit, Pieter Meysman, Bas Oldenburg, Femke van Wijk, the Dutch TWIN-IBD consortium, the Dutch Initiative on Crohn and Colitis

## Abstract

**Introduction:** The dysregulated immune-response in Crohn’s disease might result from disturbances in the T-cell receptor (TCR) repertoire. To investigate this hypothesis, we compared the peripheral TCR repertoire within T-cell subsets in twin pairs, concordant and discordant for Crohn’s disease.

**Methods:** We performed TCRα and TCRβ sequencing on peripheral flow-sorted CD4+ memory gut-homing (integrinα4β7+), non-gut-homing (integrinα4β7-), and regulatory T-cells (Tregs) from Dutch monozygotic Crohn’s disease concordant twins (N=8), monozygotic Crohn’s disease discordant twins (N=8), and healthy controls (N=4). TCR diversity and clonality, overlap between individuals, convergence and enrichment, and clustering of the TCR repertoire was studied and compared to previously reported Crohn’s disease-related TCRs.

**Results:** Overall diversity and clonality was comparable between Crohn’s disease patients, healthy cotwins and healthy controls. Comparing T-cell subsets, a decreased diversity and increased clonality was observed for Tregs. Concordant Crohn’s disease twins had an increased overlap in TCRs for Tregs and CD4+ memory gut-homing T-cells. Using TCR convergence, enrichment and subsequent clustering analyses, we identified eight clusters of TCRs potentially related with Crohn’s disease. The identified Crohn’s disease-related TCR signatures have not previously been described in relation to Crohn’s disease, and have thus far mostly unknown antigen specificity.

**Conclusions:** Increased overlap in the TCR repertoires of monozygotic twin pairs concordant for Crohn’s disease suggest that (antigen-driven) skewing of the TCR repertoire could play a role in the pathophysiology of Crohn’s disease. The identified TCR-based Crohn’s disease signatures are prime targets for further study into the pathogenesis of Crohn’s disease.

## INTRODUCTION

At the intestinal mucosa, the immune system discriminates between antigens orginating from commensal and pathogenic microbiota, balancing immune response and tolerance. In the setting of Crohn’s disease (CD) the homeostasis is disrupted leading to an impropriate and persistent inflammation.^1^ The cause of this disturbance is currently poorly understood. A diverse T-cell receptor (TCR) repertoire, consisting of a wide variety of unique TCR sequences across different T cell subsets, is needed to discriminate self from non-self and to defend the body from pathogens.^2^ In several immune-mediated diseases, e.g. rheumatoid arthritis, psoriathic arthritis, systemic lupus erythematosus, and immunodeficiency disorders, disturbances of the TCR repertoire such as differences in TCR diversity and clonal expansion have been found.^3,4^

Previous studies into the TCR repertoire in CD reported lower diversity and increased oligoclonality, i.e. presence of hyperexpanded T-cell clones, in the peripheral blood and/or intestinal mucosa of patients with CD.^5–7^ In non-responders to therapy more within-patient overlap in the mucosal TCR repertoire before and after induction therapy with adalimumab, infliximab or budesonide has been described.^5^ Postoperative recurrence after ileocecal resection has also been linked to more high-frequency mucosal T-cell clones and lower TCR diversity at the moment of surgery, and more overlap between the TCR-repertoire at the moment of surgery and 6 months thereafter.^8^ Preliminary data on the intestinal mucosal TCR repertoire in therapy-refractory CD showed a reduction in mucosal T-cell hyperexpansion following haemopoietic stem cell transplantation in responders,^9^ although another stem-cell transplantation study in CD contradicted these findings.^10^ A less diverse, more oligoclonal TCR repertoire with specific TCR-clonotypes might thus play a role in the pathogenesis of CD, suggesting shared antigenic drivers or convergent T-cell responses within CD.

The presence of dominant T-cell clones raises the question what the cognate antigen(s) for these TCRs might be. Linking newly identified antigens to these TCRs could reveal important clues regarding the underlying disease mechanisms. Indeed, efforts have been made to identify CD-associated clonotypes. The aforementioned study on ileocecal resection did not identify specific shared ileal TCR clonotypes, probably due to the high variation in TCRs between patients.^8^ Nonetheless, a different approach focusing on the TCRα and TCRβ repertoire in peripheral blood identified a TCRα CDR3 motif that was more often found in CD patients than healthy controls or ulcerative colitis patients.^11,12^ T-cells expressing these TCRα clonotypes had a gene-expression profile similar to mucosal associated invariant T-cells and natural killer T-cells, and were coined CD associated invariant T-cells (CAITs). Interestingly, TCRβ clonotypes associated with these CAITs were highly variable between patients.^11^ Via a different approach, more than 1000 TCRβ-clonotypes potentially linked to CD were recently identified by analysing the bulk TCRβ repertoire of healthy controls, CD and ulcerative colitis patients.^7^ None of these TCRβ-clonotypes overlapped with the TCRβ repertoire of the earlier described CAITs, however.^7,11^

Studying the TCR-repertoire in CD concordant en discordant monozygotic twin pairs can be a highly informative approach for identifying disease-related TCR clonotypes, because the human leukocyte antigen (HLA) is shared between monozygotic twins, as are genetic and (childhood) environmental factors. Previous studies in healthy monozygotic twins have indeed shown an increase in the shared TCR sequences in peripheral blood both for CD4+ and CD8+ T-cells.^13,14^ If a TCR sequence is shared more frequently among concordant twin pairs, this suggests a stronger association between that TCR and the disease, bringing the identification of CD-related TCR sequences within reach. An earlier study into the bulk blood TCR-repertoire in twins with inflammatory bowel diseas (IBD) including a small number of CD patients identified a few clonotypes, potentially associated with IBD, but did not specifically look at the CD-associated clonotypes.^15^ In addition, most studies have analyzed the TCR repertoire of total T-cells, whereas focusing on specific T-cell subsets may offer greater resolution in identifying CD–associated TCRs. In particular, potentially gut-homing (integrin α4β7⁺) T-cells and regulatory T-cells (Tregs) are of interest.

In this study we analyzed the peripheral TCR repertoire in a unique cohort of monozygotic twins concordant or discordant for CD, aiming to identify shared clones as well as enriched or convergent TCRs beyond simple sequence overlap. Of note, most studies in CD focused on exact TCR-clonotype matching between patients while it is known that not only exact but also similar TCR-sequences might recognize the same cognate antigen.^16^ By clustering of TCR sequences, we expect to gain better insight into CD-specific TCR motifs. To provide additional biological context, analyses were performed within defined T-cell subsets, including memory CD4+ gut homing (integrin α4β7+), memory CD4+ non-gut homing (integrin α4β7-), and CD4+ regulatory T-cells (Tregs).

## MATERIAL AND METHODS

### Study population

We included twin pairs from the ongoing prospective longitudinal Dutch “Twin cohort for the study of (pre)clinical inflammatory bowel disease in the Netherlands” (TWIN-IBD study; Dutch Trial Register: NL6187).^17^ For the present cross-sectional study, we included 1) CD concordant monozygotic twin pairs (i.e., both twins of the twin pair are affected by CD) and 2) CD discordant monozygotic twin pairs (i.e., one of the twins of the twin pair is affected by CD). Twins affected by CD from CD concordant and discordant twin pairs are referred to as “Crohn’s disease twins” and individuals who are not diagnosed with IBD from CD discordant twin pairs are referred to as “healthy cotwins”. Unrelated individuals not affected by IBD, referred to as “healthy controls”, were recruited via the Mini Donor Service at the University Medical Center (UMC) Utrecht, the Netherlands.

### Ethical considerations

The research was carried out in accordance with the declaration of Helsinki and the Dutch Medical Research Involving Human Subjects Act. The TWIN-IBD study was approved by the medical ethics committee of the UMC Utrecht (NL61114.041.17). All participants provided informed consent.

### Blood sampling, processing, and TCR-sequencing

#### Sample collection and PBMC isolation

Peripheral blood was collected from all participants in sodium heparin tubes. Peripheral blood mononuclear cells (PBMCs) were isolated using Ficoll Isopaque density gradient centrifugation (GE Healthcare Bio-Sciences, AB) and were subsequently gradually frozen at - 80°C in RPMI1640 medium containing 1% penicillin/streptomycin (Gibco), 1% L-glutamine (Thermo-Fisher Scientific, Life Technologies), 20% Fetal Calf Serum (FCS) (Invitrogen) and 10% dimethyl sulfoxide (DMSO, Sigma-Aldrich) using a Corning CoolCell container (Sigma-Aldrich). The PBMCs were thereafter stored in liquid nitrogen until further use.

#### Flow cytometry and cell sorting

PBMCs were thawed and resuspended in RPMI1640 medium containing 1% penicillin/streptomycin, 1% L-glutamine, and 10% FCS. PBMCs were first incubated with fixable viability dye eFluor506 (1:1000; eBioscience) in phosphate buffered saline (PBS) for 20 minutes at 4°C and washed in FACS buffer (cold PBS with 2% FCS and 0.1% sodium-azide (Severn Biotech Ltd.)). For surface staining, cells were subsequently incubated for 20 minutes at 4°C in FACS buffer containing 2% mouse serum (Bioconnect) and 8% Brilliant stain buffer (BD Biosciences) with the following antibodies directed against human CD3, CD4, CD8α, CD25, CD45RA, CD127, CCR7, integrin α4, integrin β7, and CCR9, and viability dye (supplementary table 1). Alive CD3+CD8-CD4+CD25+CD127low (Tregs), CD3+CD8- CD4+CD25-CCR7+/-CD45RA-Integrinα4β7+ (CD4 gut-homing memory T-cells), CD3+CD8- CD4+CD25-CCR7+/-CD45RA-Integrinα4β7- (CD4 non gut-homing memory T-cells) cells were sorted (supplementary figure 1). Between 45.000 and 100.000 cells were sorted per subset. Sorted T-cell subsets were lysed in 450μL TRIzol LS Reagent (Invitrogen) and stored at −80°C until RNA isolation.

To check for forkhead box P3 (FOXP3) expression of the sorted Treg populations, a subsample of sorted Tregs, and non-Tregs (alive CD3+CD8-CD4+CD25-) T-cells were fixed and permeabilized using 1 part fixation/permeabilization concentrate and 3 parts fixation/permeabilization diluent (eBioscience) for 30 minutes at 4°C and subsequently stained for 30 minutes at 4°C in 10x diluted Permeabilization buffer (eBioscince) containing 1% rat serum (Thermofisher) and an anti-human FOXP3 antibody (supplementary table 1 and supplementary figure 1). Sorting and data acquisition were performed on a FACSAria™ III (BD) and analyzed using FlowJo Software v10 (Tree Star Inc.).

#### TCR-sequencing

For total RNA extraction, after thawing at room temperature, 120μL chloroform was added to the vials. The vials were shaken well, incubated at room temperature and spun down at 12.000g for 15 minutes at 4°C. The aqueous phase was transferred into new vials. RNA was precipitated with 300μL isopropanol in the presence of GlycoBlue (Invitrogen) and incubated for 1 hour at -20°C. After spinning down at 12.000g for 10 minutes at 4°C the supernatant was carefully discarded and the RNA pellet was washed twice with 562.5μL 75% ethanol.

The dried pellet was then dissolved in 15μL RNAse free water and stored at -80°C until quality control and library preparation. The purity, integrity and quantity of the isolated RNA was checked with the 2100 bioanalyzer (Agilent) with pico or nano kit as appropriate and the qubit (Invitrogen) according to the manufacturers’ protocol.

Nine μL of the isolated RNA in RNAse free water was used per sample to construct TCR receptor repertoire libraries using the SMARTer Human TCR a/b Profiling Kit v2 (Takara bio), a unique molecular identifier (UMI)-based 5’ rapid amplification of complementary DNA (cDNA) ends (5’ RACE)-like method using switching mechanism at 5’ end of RNA Template (SMART) full-length cDNA synthesis technology, according to the manufacturer’s protocol.

By incorporating UMIs, we were able to correct for potential polymerase chain reaction (PCR) bias after sequencing. In short, reverse transcription was performed with a TCR dT primer and further extended with non-templated nucleotides, to which the TCR SMART UMI oligo anneal, using MMLV derived SMARTScribe^TM^ reverse transcriptase. Next, the cDNA was amplified in two semi-nested PCR-steps. In the first PCR-step TRA and TRB constant regions reverse primers and a human TCR universal forward primer were used. In the second PCR-step the PCR1 amplicons were targeted with reverse and forward primers including adapter and unique dual index (UDI) sequences, enabling sample barcoding.

NucleoMag Next-Generation Sequencing Clean-up and Size Select beads were used to purify the PCR products.

Before sequencing, pools were made containing libraries of 24 samples per pool, and each pool was purified aiming to enrich for 200-1000bp long cDNA. The pools were then sequenced twice on an Illumina NovaSeq6000 platform (Genomescan, the Netherlands) to generate at least 10 million 150 base pairs long, paired-end, reads per sample.

#### TCR sequencing data processing

The quality of the sequence results was checked with the multiQC toolkit. FASTQ raw data files were processed with Cogent NGS Immune Profiler Software version 1.0 (Takara Bio) which incorporates MIGEC version 1.2.9 and MiXCR version 2.1.8.^18,19^ We set identical molecular identifier group (MIG) thresholds for all samples: 6 for TCRα and 3 for TCRβ.

#### Analyses

MiXCR output was combined across all patients and non-productive TCRs, containing non-standard amino acids or unproductive V- and J-genes, were removed. Unique clonotypes are defined as a unique combination of V-, (D-) and J-fragments per patient and per cell type.

Unique clonotypes were used to determine repertoire richness as well as Pielou evenness, which is a normalized Shannon diversity, using scikit-bio alpha diversity (v0.5.9, Python v3.8.18). Richness and Pielou evenness were determined per chain type (TCRα and TCRβ) and patient group (Crohn’s disease twins, healthy cotwins, healthy controls). Exact overlap as well as the adjusted Morisita-Horn index were used to quantify repertoire overlap between concordant twins, discordant twins, unrelated CD pairs (i.e. Crohn’s disease twins matched to an unrelated CD patient) and unrelated healthy pairs (i.e. healthy cotwins and healthy controls). In contrast to the original Morisita-Horn index, this adjusted version emphasizes both on the presence and relative frequency of shared clonotypes, and reduces the effect of private non-overlapping clones, using the following formula implemented in Python:

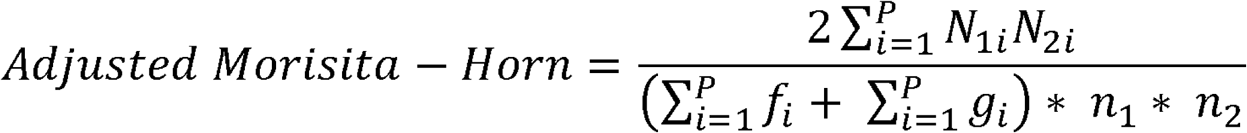

With P representing all shared, public clonotypes and N_1i_ and N_2i_ representing the absolute abundance of shared clone i in both repertoires. The clonal relative frequency in both repertoires is given as f_i_ and g_i_, summed across all shared clones. Finally, n1 and n2 represent the total abundance per repertoire. Differences in overlap between the 4 groups was calculated using the Kruskal-Wallis test, followed up by posthoc pairwise Dunn’s test if there was a significant group difference and Bonferroni multiple testing correction.

TRIASSIC ^20^ [v0.2.0, Python v3.10.14] was used to identify TCRs that were more convergent in the CD group, compared to healthy cotwins and healthy controls. TRIASSIC compares clonotypes across the full cohort and quantifies convergence of a TCR by counting clonotypes that are similar, but developed independently. These events of convergent recombination were identified by the presence of (i) the same clonotype in a different individual, (ii) a clonotype with differing nucleotide sequence coding for the same amino acid sequence regardless of individual and (iii) a highly similar clonotype (TCRdist^16,21^ score ≤ 12.5), regardless of the individual. The TCRdist threshold corresponds with approximately one amino acid sequence difference in TCR sequence, allowing for the identification of TCR that potentially recognize the same antigen and thereby providing biologically meaningful results. For each TCR, we identified the number of convergence events within its 2048 most similar clonotypes, as identified by TCRdist. A two-tailed Fisher’s exact test is used to calculate enrichment in the number of convergent events originating from one group or another. Clonotypes that were statistically significantly associated with CD (enrichment > 0, p-value ≤ 0.05) were selected. To reduce potential effects of pre-existing/preclinical CD patterns within the healthy cotwins (who are at increased risk of developing CD), convergent TCRs that were only found when comparing against the healthy cotwin group and not healthy controls were excluded. Clonotypes that are significantly convergent in CD were then clustered based on sequence similarity (Hamming distance=1) using ClusTCR ^22^ (v1.0.2) across the full cohort. Features of these CD convergent clusters, such as sample distribution, clone counts, cell type distribution and epitope annotation, were extracted for the most promising clusters. Epitope specificity per clonotype was predicted using the ImmuneWatch Detect tool ^23^ at the default confidence level 0.2.

Significantly neighbor enriched (SNE) TCRs (defined by TCRdist unit radius ≤ 18.5, Valkiers et al. manuscript in preparation) were identified using a synthetic TCR background (10x size of original repertoire). The TCRdist unit radius is recommended to be set at 18.5 to enable to detect TCRs with one to two amino acid sequences differences, enabling to capture local clusters within an individual’s TCR repertoire. This was generated by random breaking, shuffling and re-pairing of VDJ fragments of the clonotypes within the repertoire. Based on the background, expected neighbor counts were calculated and compared to the actual observed neighbors per repertoire and per chain type. Enrichment of these SNE TCRs was calculated using SciPy’s (v1.8.0) hypergeometric survival function and adjusted for multiple testing using the Bonferroni correction. Further prioritization was applied by selecting those TCR clusters that were significantly convergent for CD and that contained TCR sequences that were SNE TCRs.

Lastly, we compared all unique TCRs in the present study to TCRs potentially associated with CD as reported in the literature. To this end, we compared all the unique TCRs, the unique TCRs within the CD patients specifically and the TCRs identified in our clustering analysis to the TCRs identified in literature. This includes the study of Rosati et al. 2020^15^ in twins concordant and discordant for IBD, the study of Rosati et al. 2022^11^ which identified CAITs in their dataset, and the study of Rios Martini et al 2023^24^ which reported an extensive list of yeast-reactive TCRs in the setting of CD. Finally, we aimed to compare against the “enhanced TCR sequences” associated with CD as described by Pesesky et al. 2025^7^, unfortunately the TCR sequences could not be shared by the authors, and therefore we compared to a patent^25^ based on earlier analyses of the same group, which includes 2010 unique TCRs.

## RESULTS

### Baseline characteristics

TCR-sequencing was performed for 4 CD concordant monozygotic twin pairs (8 individuals), 4 CD discordant monozygotic twin pairs (8 individuals), and 4 healthy controls (table 1, figure 1 and supplementary figure 2). TCR-repertoire data was generated for all 20 individuals for Tregs and CD4 gut-homing memory T-cells, and for 1 CD concordant twin pair, 2 CD discordant twin pairs, and 2 healthy controls for CD4 non-gut-homing memory T-cells.

**Figure 1.**
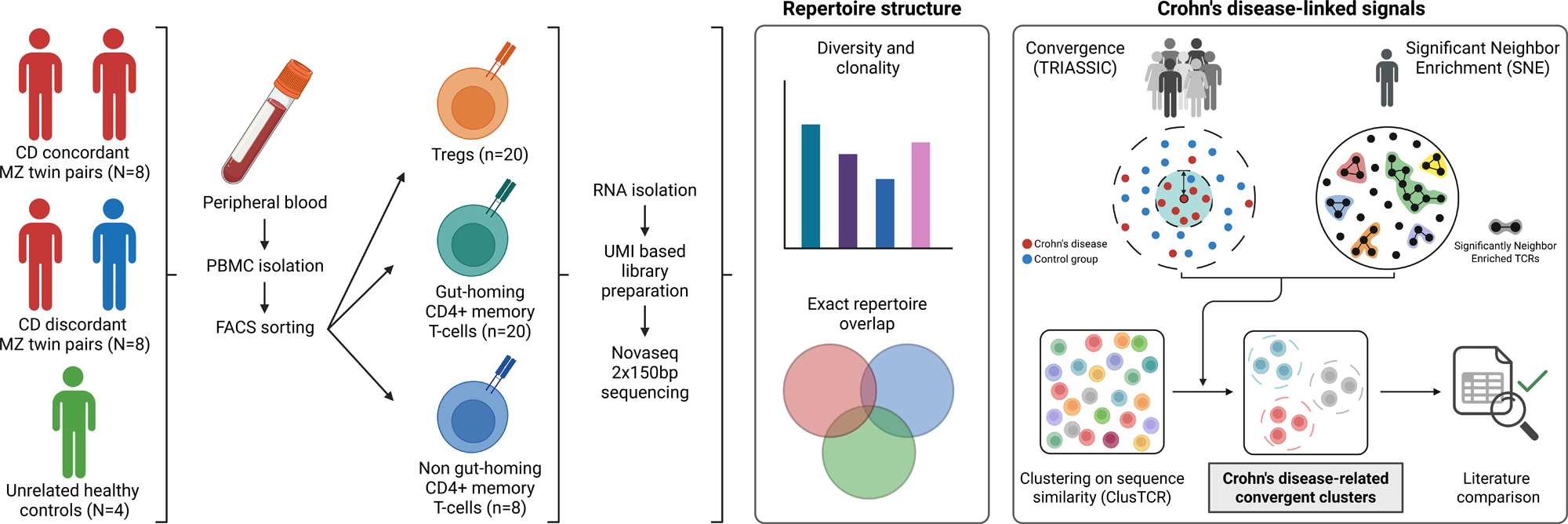
Study overview. From PBMCs of 4 monozygotic twin pairs concordant for Crohn’s disease, 4 monozygotic twin pairs discordant for Crohn’s disease, and 4 unrelated healthy controls regulatory T-cells (alive CD3+CD4+CD25+CD127-), gut-homing CD4+ memory (alive CD3+CD8-CD4+CD25-CCR7+/-CD45RA-integrinα4β7+) T-cells, and non-gut homing CD4+ memory (alive CD3+CD8-CD4+CD25-CCR7+/-CD45RA-integrinα4β7-) T-cells were FACS-sorted (supplementary figure 1). RNA was isolated and used to prepare a UMI-based TCR-library for 2x150bp sequencing on a Novaseq platform. From this, TCRα and TCRβ repertoire analyses were performed. Diversity, clonality and repertoire overlap was studied. Subsequently, TCR convergence analyses followed by clustering, and TCR significant neighbor enrichment analyses followed by clustering were performed. Both methods identify not only exact amino acid matches but also similar sequence matches against a TCR background, across the whole cohort or within a single repertoire respectively. As a final step we combined the results of both analyses to identify the most promising CD-related TCR cluster, and compared the identified CD-related clusters to the existing literature. bp, basepairs, CD, Crohn’s disease; FACS, fluorescence-activated cell sorting; gut-homing CD4+ memory T-cells, alive CD3+CD8-CD4+CD25-CCR7+/-CD45RA-Integrinα4β7+ T-cells; MZ, monozygotic; N, number of participants; n, number of samples; non gut- homing CD4+ memory T-cells, alive CD3+CD8-CD4+CD25-CCR7+/-CD45RA- Integrin-α4β7- T-cells; PBMC, peripheral blood mononuclear cells; TCR, T-cell receptor; Tregs, regulatory T-cells, alive CD3+CD4+CD8-CD25+CD127low T-cells; UMI, unique molecular identifier.

**Table 1.**
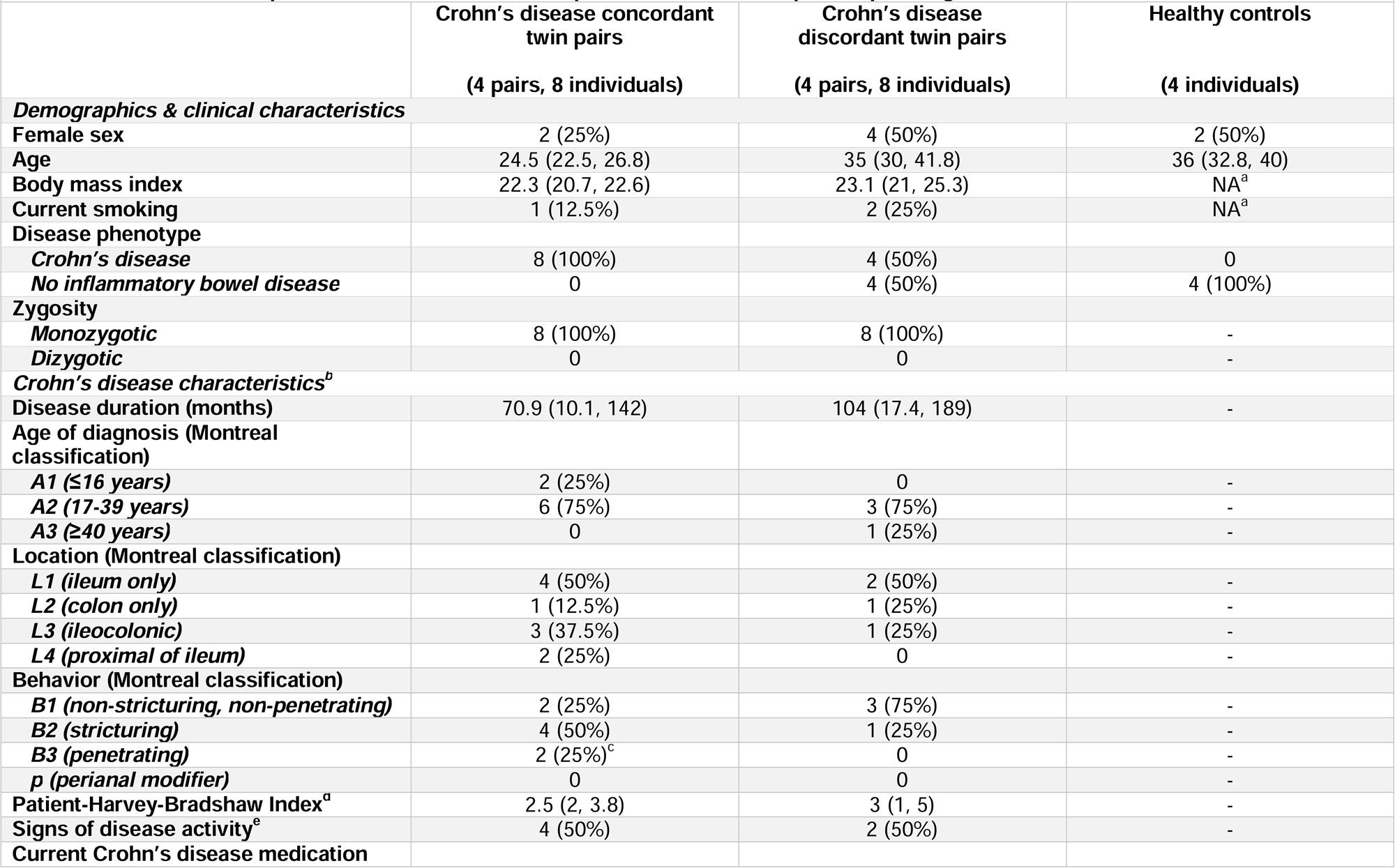
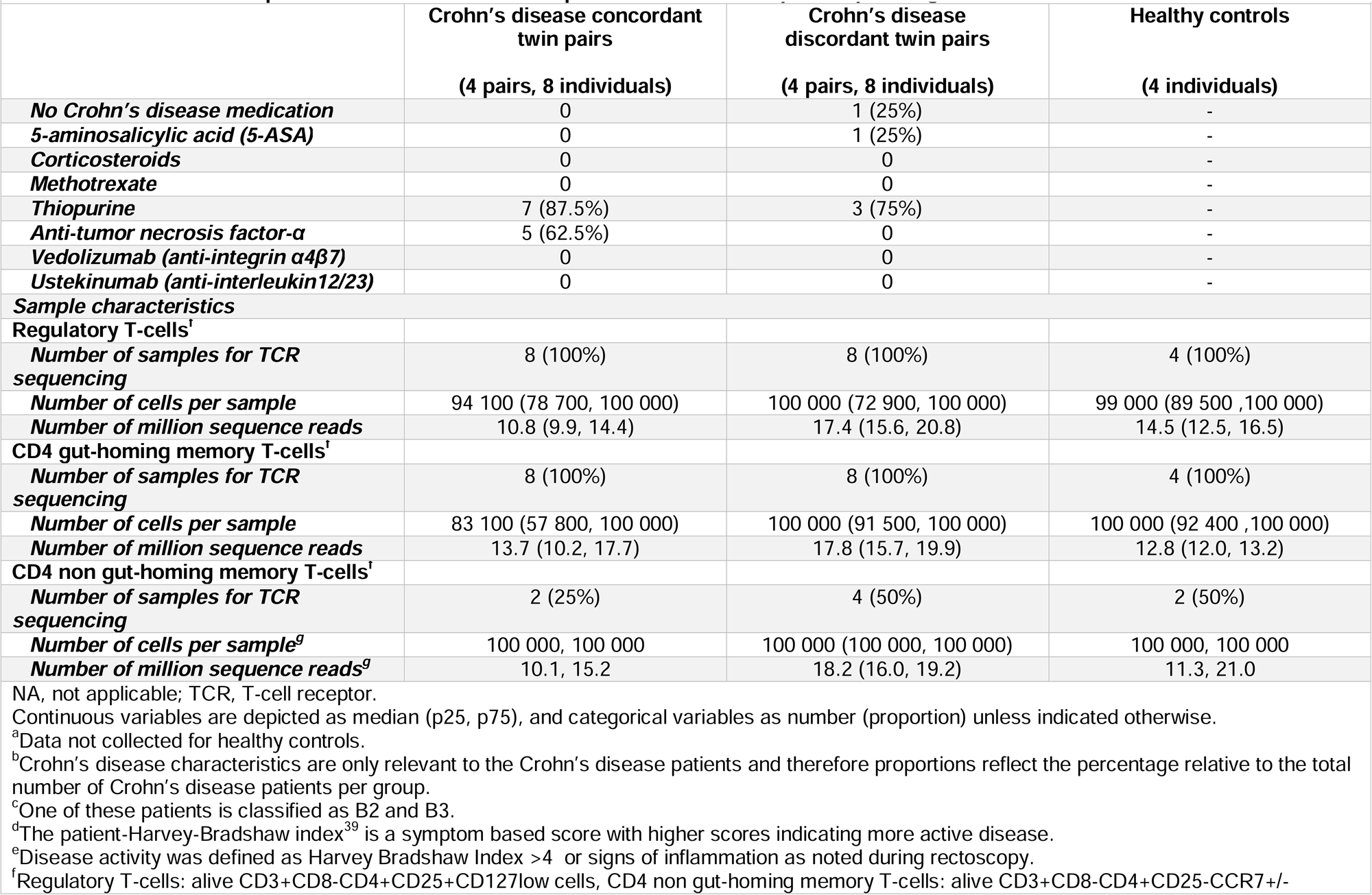
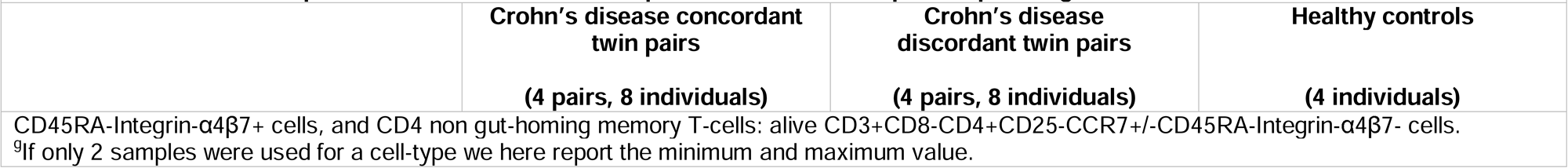
Baseline and sample characteristics of included patients for T cell receptor sequencing.

Concordant CD twin pairs were younger than discordant CD twin pairs and healthy controls (median age 24.5, 35, and 36, respectively). The CD patients within the concordant twin pairs compared to the CD patients from discordant twin pairs, were often younger at diagnosis (Montreal classification A1: 25% vs. 0% respectively), and had more often stricturing and penetrating disease (Montreal classification B2: 50% vs. 25%, B3: 25% vs. 0% respectively). The concordant and discordant CD twins were comparable regarding patient Harvey-Bradshaw Index (median: 2.5 vs. 3), proportion of patients with signs of disease activity (50% vs. 50%), and use of thiopurines (87.5% vs 75%), although only CD patients from concordant twin pairs used anti-tumor necrosis factor-α (62.5%).

No statistically significant difference in the proportion of Tregs, gut-homing CD4+ memory T-cells, or non-gut-homing CD4+ memory T-cells was found comparing the 16 CD twins, 4 healthy cotwins and 6 healthy controls for whom flow-cytometry data was available (supplementary table 2 and supplementary figure 3).

We generated a median of 15.1 (p25-p75: 12.1-18.5) million sequence reads per sample containing median 54.8 (p25-p75: 49.7, 56.9) percentage TCRα, median 43.9 (p25-p75: 41.9, 48.7) TCRβ, and median 1.1 (p25-p75: 0.9-1.5) undetermined reads. After UMI-correction a median number of 13400 (p25-p75: 6530-24800) TCRα and 20400 (p25-p75: 11300-40700) TCRβ clonotypes remained.

#### Richness, diversity and clonal expansion

No clear difference was found in richness (i.e. unique TCR sequences), diversity (Pielou’s evenness), and clonal expansion in the TCRα and TCRβ repertoire between CD, healthy cotwins and healthy controls (Figure 2). The diversity in the TCRα and TCRβ repertoires of Tregs was numerically lower compared to the TCR repertoires of gut-homing CD4+ memory T-cells and non-gut-homing CD4+ memory T-cells, irrespective of disease state. In line with this, the TCRβ repertoire, and to a lesser extent the TCRα repertoire from Tregs was more clonally expanded (Figure 2).

**Figure 2.**
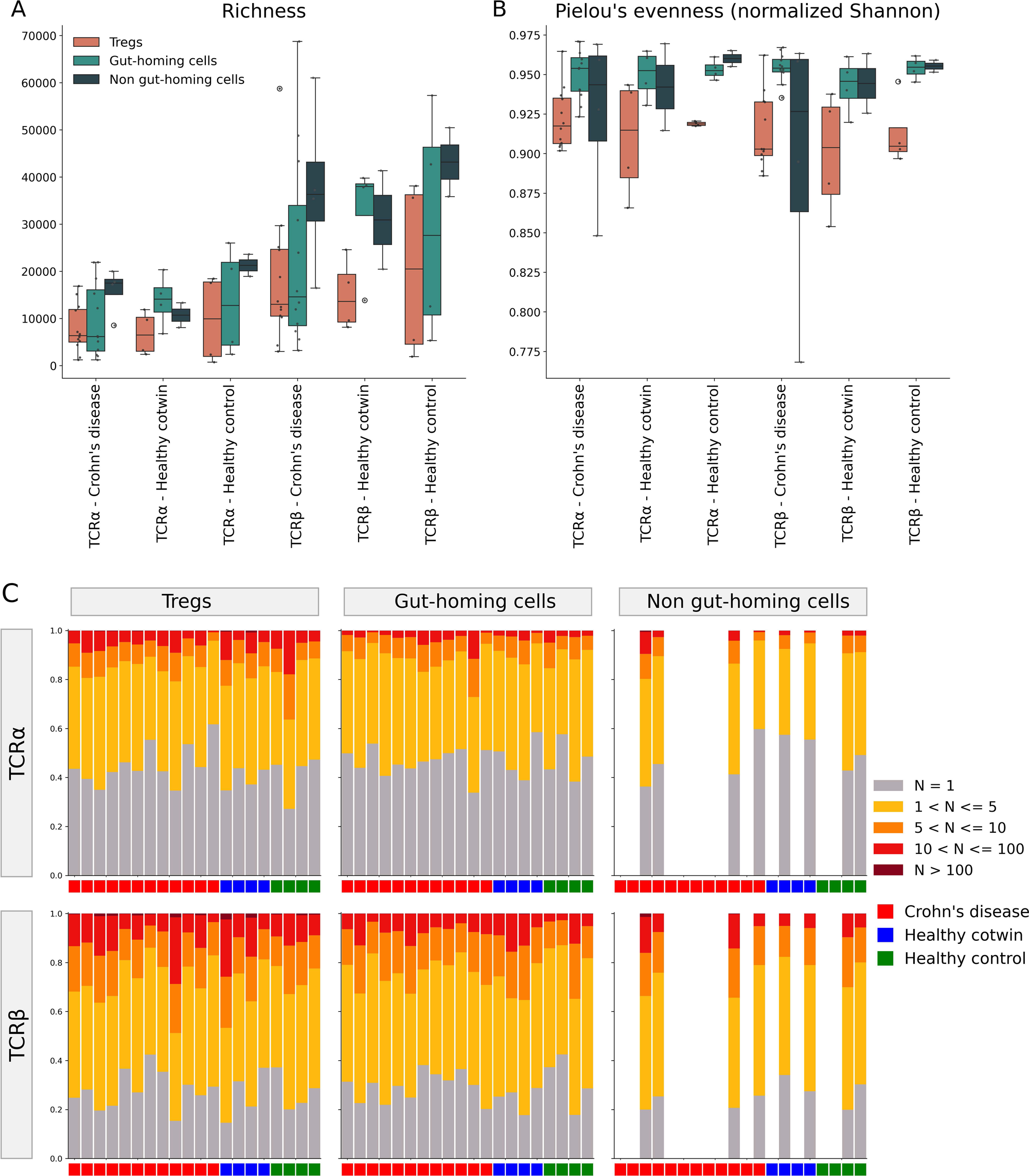
Richness, diversity and clonal expansion. A) richness (i.e. the number of unique TCR-sequences) and B) diversity (expressed as Pielou evenness, which is a normalized Shannon metric) are displayed per cell type, separately for TCRα and TCRβ sequences, subdivided for Crohn’s disease patients, healthy cotwins and unrelated healthy controls. Overall the richness and diversity is lower for Tregs compared to gut-homing and non-gut-homing CD4+ memory T-cells, irrespective of disease subtype. C) absolute clonal expansion (i.e. the number of T-cells per unique TCR) per cell type, separately for TCRα and TCRβ sequences is displayed per individual. Especially in the TCRβ sequences clonal expansion of the Tregs is noted. gut-homing cells, alive CD3+CD8-CD4+CD25-CCR7+/-CD45RA-Integrinα4β7+ T- cells; N, number of T-cells; non gut-homing CD4+ memory T-cells, alive CD3+CD8- CD4+CD25-CCR7+/-CD45RA-Integrin-α4β7- T-cells; TCR, T-cell receptor; Tregs, regulatory T-cells, alive CD3+CD4+CD8-CD25+CD127low T-cells.

#### Overlap in TCR sequences

We hypothesized that if the TCR repertoire is associated with CD, a higher overlap within concordant CD twin pairs could be expected compared to discordant CD twin pairs and pairs of unrelated individuals. For TCRβ sequences, a statistically significant (Bonferroni adjusted p ≤ 0.05) higher overlap (expressed as the adjusted Morisita-Horn index, taking size of the different repertoires into account) was noted within the concordant CD pairs compared to pairs of unrelated healthy individuals (i.e. healthy cotwins and healthy controls) for both Tregs and gut-homing CD4+ memory T-cells (Figure 3). Overall, there was a numerical trend toward lower overlap within discordant CD twin pairs. No overlap analyses were performed for the non-gut-homing CD4+ memory T-cells, because TCR repertoires for these cell types were only determined in a subgroup of participants. For the TCRα repertoire only for Tregs an increase in the adjusted Morisita-Horn index comparing concordant CD twin pairs to unrelated pairs of healthy individuals was found. These results suggest that based on overlap analyses there is skewing of the peripheral TCRβ repertoire of Tregs and gut-homing CD4+ memory T-cells associated with CD.

**Figure 3.**
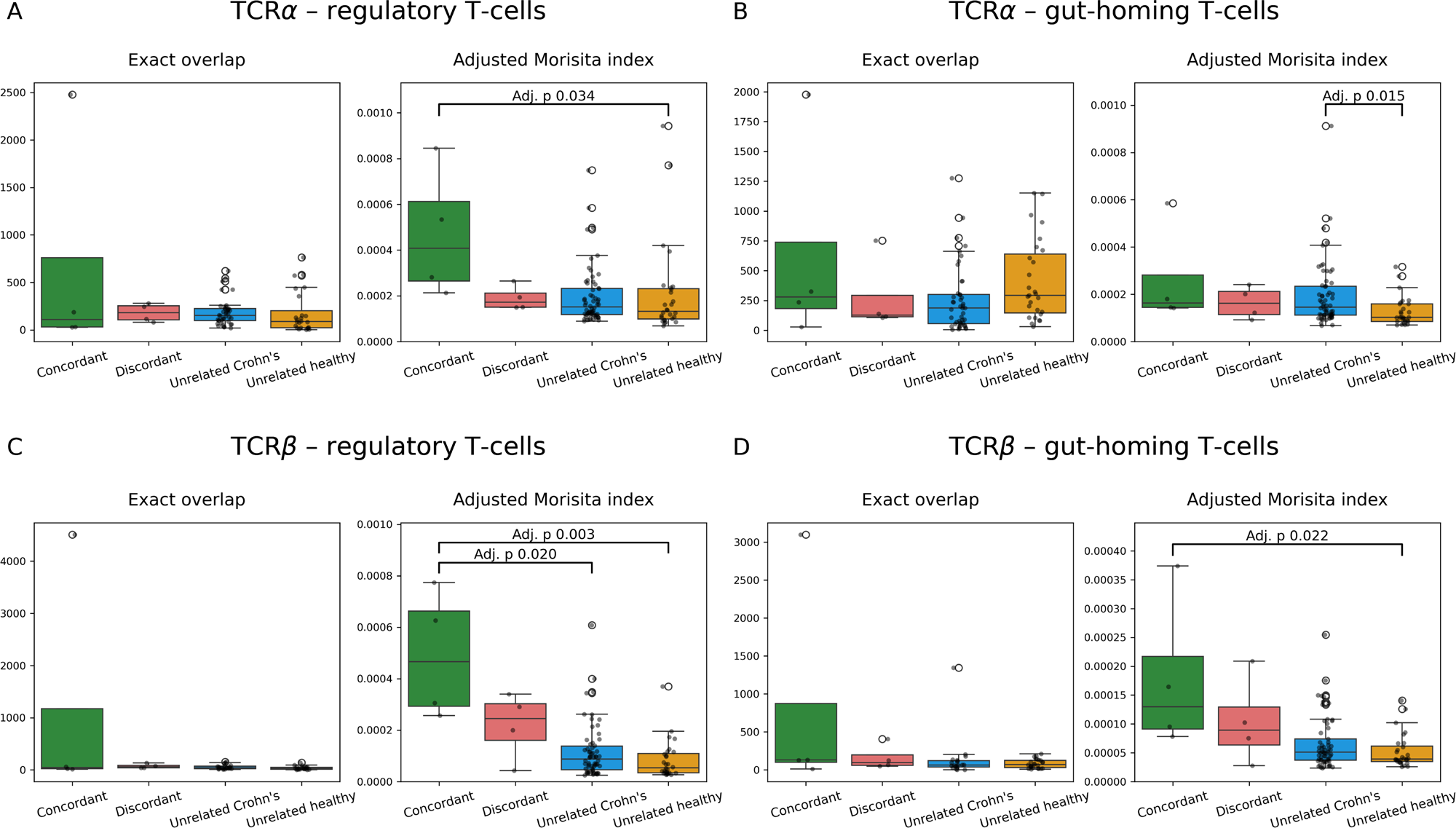
Overlap between individuals in TCR-sequences. Exact clonotype overlap and the adjusted Morisita-Horn index is displayed within Crohn’s disease concordant twin pairs, Crohn’s disease discordant twin pairs, pairs of unrelated Crohn’s disease patients, and pairs of unrelated healthy participants (i.e. healthy cotwins and healthy controls). Results are separately displayed for TCRα (A and B), and TCRβ (C and D) for regulatory T-cells (A and C) and gut-homing CD4+ memory T-cells (B and D). There is a statistically significantly (Bonferroni adjusted p ≤ 0.05) increased overlap within the TCR repertoire within concordant Crohn’s disease twin pairs compared to unrelated healthy controls for regulatory T-cells and gut-homing CD4+ memory T-cells within the TCRβ repertoire, and for regulatory T-cells in the TCRα repertoire. Gut-homing CD4+ memory T-cells, alive CD3+CD8-CD4+CD25-CCR7+/-CD45RA- Integrinα4β7+ T-cells; non gut-homing CD4+ memory T-cells, alive CD3+CD8- CD4+CD25-CCR7+/-CD45RA-Integrin-α4β7- T-cells; TCR, T-cell receptor; regulatory T- cells, alive CD3+CD4+CD8-CD25+CD127low T-cells.

#### TCR convergence, enrichment, clustering, and comparison to existing literature

To further explore which TCRs might be associated with CD we took two complementary approaches, i.e. TCR convergence analyses followed by clustering, and TCR neighbor enrichment analyses followed by clustering. Both methods identify not only exact amino acid matches but also similar sequence matches against a TCR background, across the whole cohort or within a single repertoire respectively. As a final step we combined the results of both analyses to identify the most promising CD-related TCR clusters.

In TCR convergence analyses, for each TCR the number of convergence events, i.e. the same or similar clonotypes that developed independently either in a different individual, a different nucleotide sequence, or within a TCRdist distance ≤ 12.5, is calculated and compared between 1) CD patients, 2) healthy cotwins and 3) healthy controls. TCRs that showed a higher number of convergence events in CD patients compared to healthy samples, potentially indicating a shared driver for convergence, were identified using enrichment testing based on the Fisher’s exact test. The analysis revealed 8123 convergent TCRs when comparing the CD patients to all healthy individuals. A larger number of convergent TCRs was found in the CD group than within the healthy cotwins and healthy controls, suggesting a potential CD related skewing towards higher convergence in these samples. Among the convergent TCRs there was no preference for a specific cell type (Figure 4A-B, supplementary figure 4), and for most of these TCRs Immunewatch Detect was unable to predict their epitope specificity against known microbial, viral or self-antigens (Figure 4C). In order to identify groups of similar TCRs found across different CD patients, the significantly convergent TCRs were clustered based on sequence similarity using ClusTCR. This resulted in a large number of potentially CD-related clusters that varied in size and patient distribution. Each cluster now contained multiple distinct but CD-convergent TCRs, grouped based on potential recognition of shared antigens that might play a role in CD.

**Figure 4.**
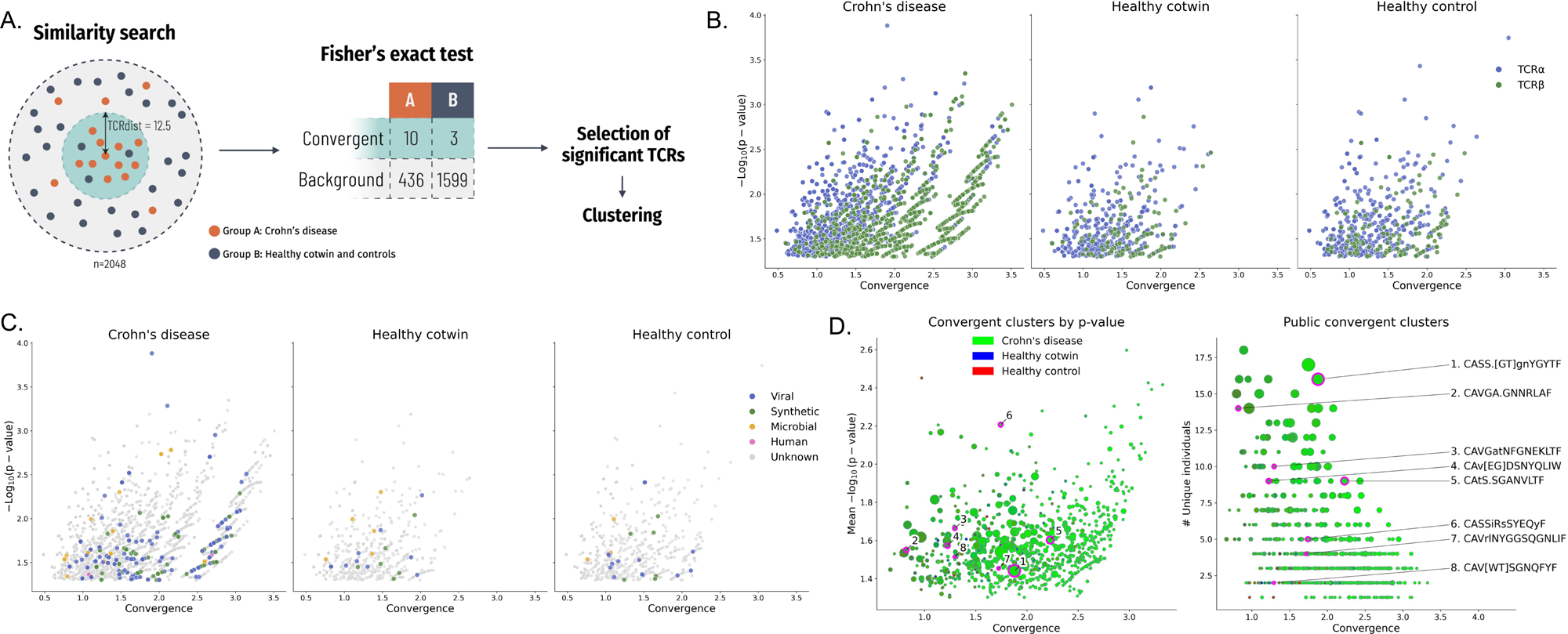
TCR convergence and potentially CD-related TCR clusters. A) schematic representation how TCR-convergence is calculated. Within a group (e.g. Crohn’s disease patients) TCRs which are similar based on a TCRdist of 12.5 are counted and compared to the number of TCRs found in the comparator groups. B-C) the statistically significant (p ≤ 0.05) TCRs and their convergence measure are displayed separately for Crohn’s disease, healthy cotwins and healthy controls. The TCRs are colored by B) TCRα and TCRβ sequences and C) the predicted epitope recognition based on Immunewatch Detect^23^. The TCR convergence in general is larger for TCRβ sequences and is more pronounced in Crohn’s disease patients (B). The vast majority does not have a prediction on the epitope- specificity, with potential viral antigens being the largest group of assigned potential targets of the TCRs (C). D) The TCRs with statistically significant convergence within the Crohn’s disease group were clustered based on sequence similarity using ClusTCR. This shows multiple clusters that have increased convergence in more than one Crohn’s disease sample, left on p-value, right on number of Crohn’s disease samples with that cluster. Taking statistical significance, number of participants in whom a TCR related to a clusters is found into account combined with the TCR neighbor enrichment analyses (figure 5 and supplementary figure 5-6) eight potentially CD-related clusters (table 2 and supplementary table 3) could be identified and are highlighted in the plot. TCR, T-cell receptor.

**Table 2.**
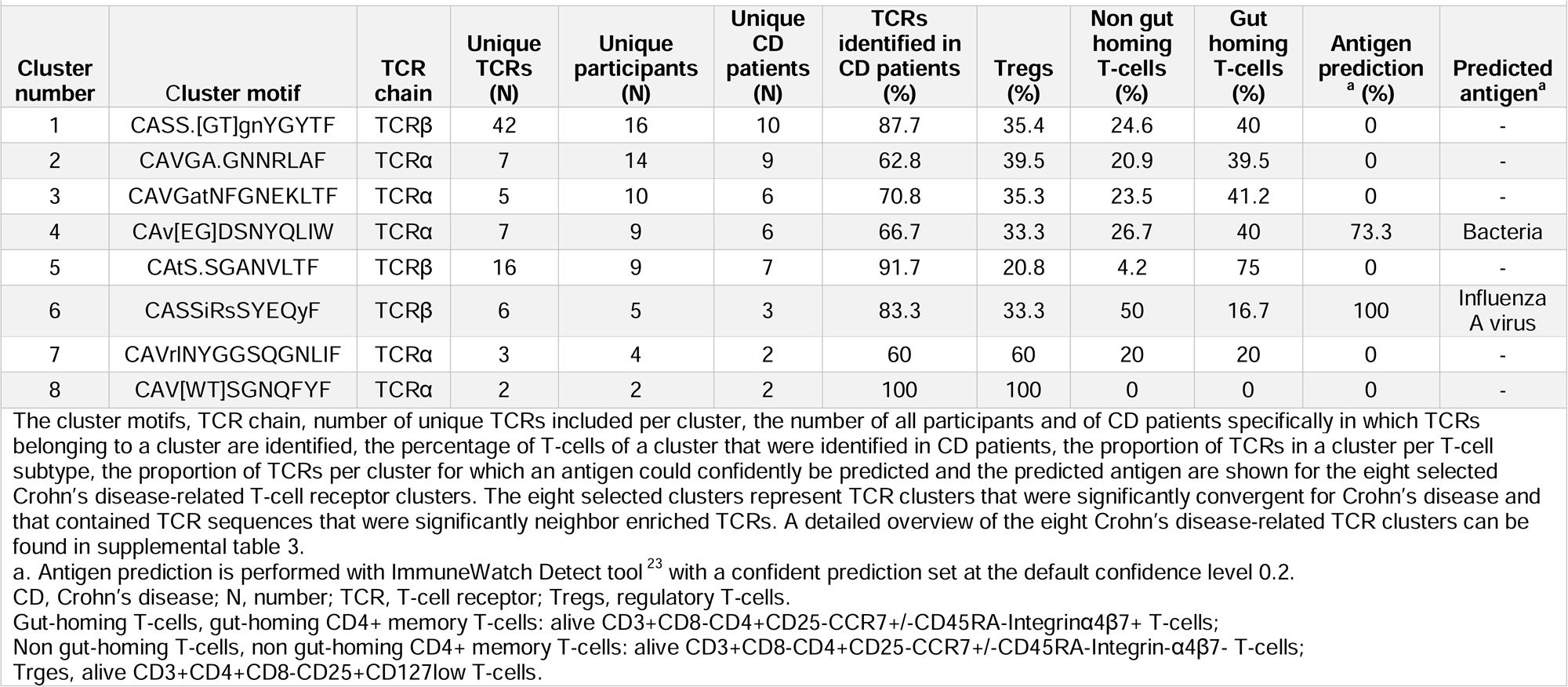
Overview of the eight selected Crohn’s disease-related T-cell receptor clusters.

To further explore the relevant convergent clusters, we first performed TCR enrichment analyses using significantly enriched neighbors In this analysis, the number of neighbor TCR sequences (TCRdist distance ≤ 18.5) within the repertoire of one individual is compared to a synthetic background. This allows for the identification of TCRs that have more similar TCRs than expected by chance within a single repertoire.

These analyses revealed different TCRs that were more enriched within individual CD patients, than within healthy cotwins and healthy controls. The most statistically significant TCRs for CD patients were TCRα sequences predicted to be directed against microbiota, likely mucosa-associated invariant T cells (MAIT cells) based on their V- and J-gene usage (TRAV1-2 combined with TRAJ12, TRAJ20 or TRAJ33). Additionally, TCRβ sequences predicted to be directed against the influenza A virus were also enriched in CD patients (figure 5A and supplementary figure 5A-C). Interestingly the most statistically significantly enriched TCRs were found in the repertoire of one individual (supplementary figure 5D).

**Figure 5.**
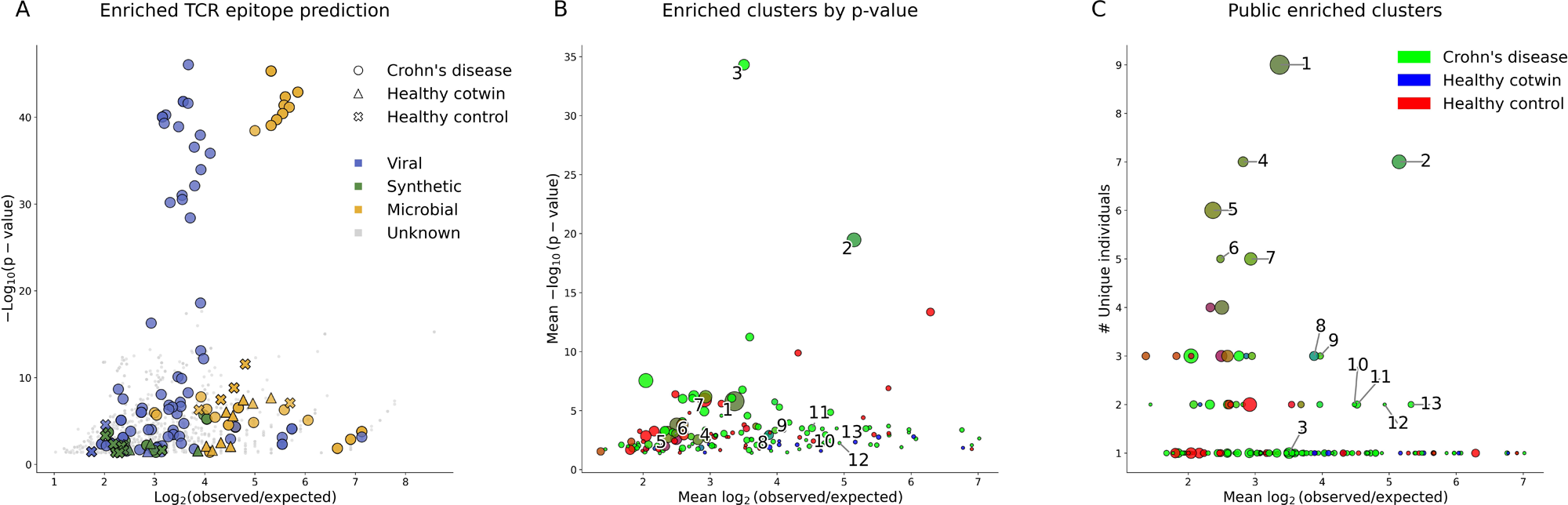
TCR neighbor enrichment analyses and clustering more detailed. TCR neighbor enrichment is calculated comparing the number of neighbor TCR sequences (i.e. similar TCR sequences) within one TCR repertoire compared to a synthetic background repertoire. A) Only statistically significantly (Bonferroni adjusted P-value ≤ 0.05) enriched TCRs are displayed. Shapes refer to whether the TCR is found in Crohn’s disease patients, healthy cotwins or healthy controls. Colors show the predicted epitope based on Immunewatch Detect^23^. Within Crohn’s disease patients a few TCRs of non-gut homing CD4+ memory T-cells which are partly predicted to be directed against viral and partly against microbial antigens are the most statistically significantly enriched TCRs. The results are displayed in more detail for TCRα and TCRβ, cell type, antigen prediction and individual participant subdivided for Crohn’s disease, healthy cotwins and healthy controls in supplementary figures 5 and 6. B and C) Clustering based on sequence similarity (ClusTCR^22^, Hamming distance = 1) was performed on the neighbor enriched TCRs. These cluster plots display the mean enrichment score, mean p-value and the number of unique patients in which TCRs from a cluster are encountered. The color gradient indicates the proportion of TCRs found across the different types of participants. Based on either exhibiting a highly significant p-value or high convergence across individuals, 13 promising clusters were selected that are mainly encountered in Crohn’s disease patients (supplementary table 3). TCR, T-cell receptor.

Subsequently, these enriched TCRs were also clustered based on sequence similarity across the different samples, reflecting cohort-wide enrichment patterns. This resulted in a total of 13 enriched clusters that seem to be of interest due to their enrichment in mainly CD patients and clusters being shared among multiple individuals. (figure 5B-C)

These 13 neighbor-enriched clusters were then used to further prioritize the list of convergent clusters from the previous analysis. Specifically, it was determined which CD-convergent clusters also contained TCRs present in any of the 13 neighbor-enriched clusters.

Convergent clusters that showed this dual signal, both convergence and neighbor enrichment, were considered as the most promising CD-related clusters. This integrative approach resulted in 8 prioritized convergent clusters (table 2, supplementary table 3 and supplementary figure 6), spanning both TCRα and TCRβ chains, which may represent CD-associated TCR patterns. It remains to be determined towards which antigens the majority of these TCRs are directed.

#### Comparison to the literature

In order to validate earlier reported potentially CD-related TCRs from literature we compared these TCRs to (1) all TCRs in our cohort, (2) all TCRs within the CD patients in our cohort, (3) all convergent clusters, and (4) the selection of 8 CD-related clusters from the present study (table 3). These comparisons were made against the suggested CD-related TCRs as reported by Rosati et al. 2020^15^ and 2022^11^, Rios Martini et al 2023^24^, and a published patent^25^.

**Table 3.**
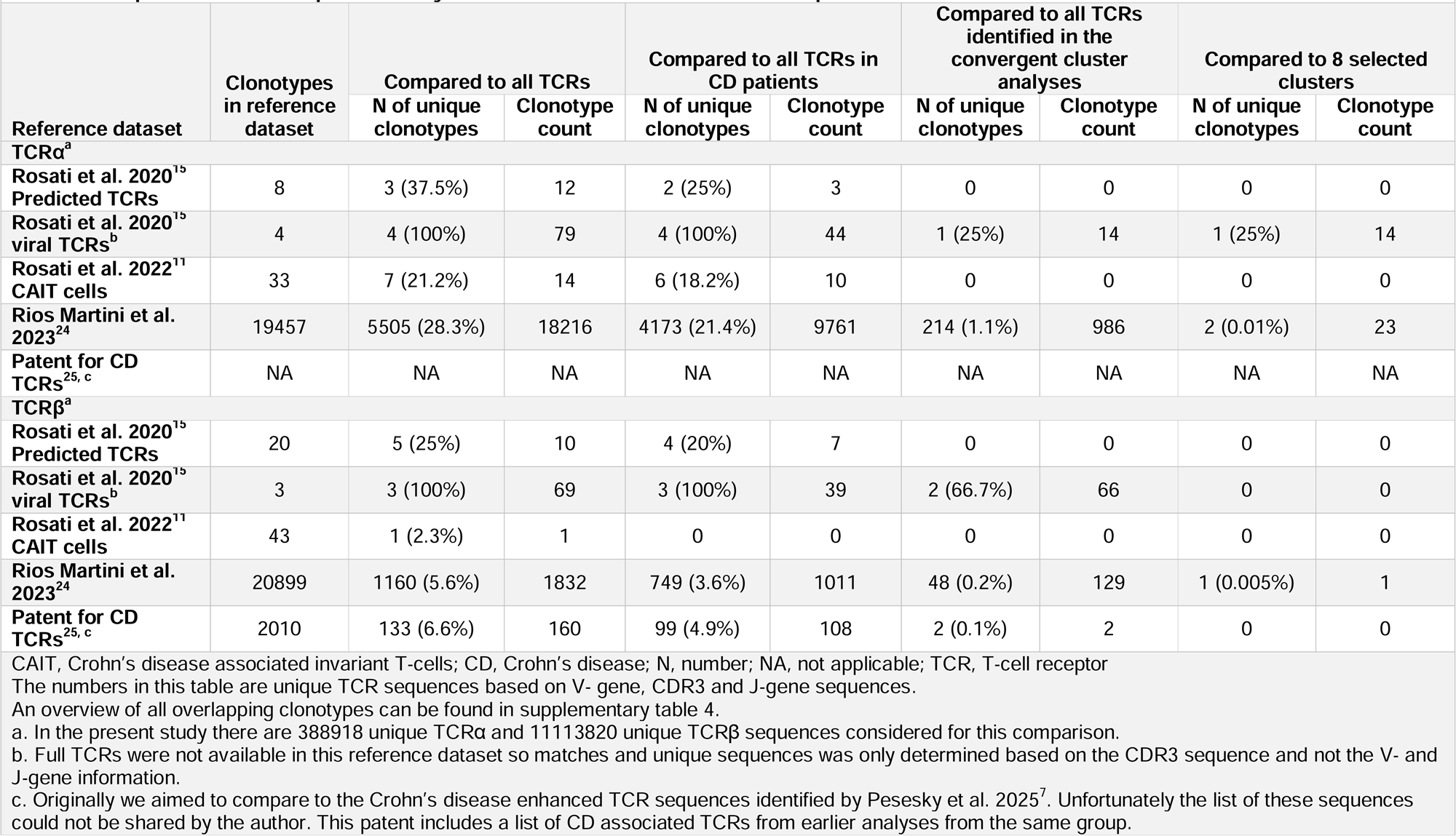
Comparison of TCRs in present study to Crohn’s disease associated TCR sequences identified in literature.

| Reference dataset | Clonotypes in reference dataset | Compared to all TCRs |  | Compared to all TCRs in CD patients |  | Compared to all TCRs identified in the convergent cluster analyses |  | Compared to 8 selected clusters |  |
| --- | --- | --- | --- | --- | --- | --- | --- | --- | --- |
|  |  | N of unique clonotypes | Clonotype count | N of unique clonotypes | Clonotype count | N of unique clonotypes | Clonotype count | N of unique clonotypes | Clonotype count |
| TCR$\alpha^a$ | | | | | | | | | |
| Rosati et al. 2020<sup>15</sup><br>Predicted TCRs | 8 | 3 (37.5%) | 12 | 2 (25%) | 3 | 0 | 0 | 0 | 0 |
| Rosati et al. 2020<sup>15</sup><br>viral TCRs<sup>b</sup> | 4 | 4 (100%) | 79 | 4 (100%) | 44 | 1 (25%) | 14 | 1 (25%) | 14 |
| Rosati et al. 2022<sup>11</sup><br>CAIT cells | 33 | 7 (21.2%) | 14 | 6 (18.2%) | 10 | 0 | 0 | 0 | 0 |
| Rios Martini et al. 2023<sup>24</sup> | 19457 | 5505 (28.3%) | 18216 | 4173 (21.4%) | 9761 | 214 (1.1%) | 986 | 2 (0.01%) | 23 |
| Patent for CD TCRs<sup>25, c</sup> | NA | NA | NA | NA | NA | NA | NA | NA | NA |
| TCR$\beta^a$ | | | | | | | | | |
| Rosati et al. 2020<sup>15</sup><br>Predicted TCRs | 20 | 5 (25%) | 10 | 4 (20%) | 7 | 0 | 0 | 0 | 0 |
| Rosati et al. 2020<sup>15</sup><br>viral TCRs<sup>b</sup> | 3 | 3 (100%) | 69 | 3 (100%) | 39 | 2 (66.7%) | 66 | 0 | 0 |
| Rosati et al. 2022<sup>11</sup><br>CAIT cells | 43 | 1 (2.3%) | 1 | 0 | 0 | 0 | 0 | 0 | 0 |
| Rios Martini et al. 2023<sup>24</sup> | 20899 | 1160 (5.6%) | 1832 | 749 (3.6%) | 1011 | 48 (0.2%) | 129 | 1 (0.005%) | 1 |
| Patent for CD TCRs<sup>25, c</sup> | 2010 | 133 (6.6%) | 160 | 99 (4.9%) | 108 | 2 (0.1%) | 2 | 0 | 0 |

CAIT, Crohn's disease associated invariant T-cells; CD, Crohn's disease; N, number; NA, not applicable; TCR, T-cell receptor
The numbers in this table are unique TCR sequences based on V- gene, CDR3 and J-gene sequences.
An overview of all overlapping clonotypes can be found in supplementary table 4.
a. In the present study there are 388918 unique TCRα and 11113820 unique TCRβ sequences considered for this comparison.
b. Full TCRs were not available in this reference dataset so matches and unique sequences was only determined based on the CDR3 sequence and not the V- and J-gene information.
c. Originally we aimed to compare to the Crohn's disease enhanced TCR sequences identified by Pesesky et al. 2025<sup>7</sup>. Unfortunately the list of these sequences could not be shared by the author. This patent includes a list of CD associated TCRs from earlier analyses from the same group.

Of the TCR sequences reported in literature, 21.2 to 37.5% of the TCRα sequences and 2.3 to 25% for the TCRβ sequences were also present in our cohort (table 3 and supplementary table 4). When only comparing the TCR sequences from literature to the TCR sequences in CD patients in our cohort the overlap was slightly lower (TCRα: 18.2 to 25%, TCRβ: 2.9 to 20%). When looking at our convergent cluster analyses, none of the reported CAIT TCR sequences^11^ were found. A small proportion of the reported yeast-reactive TCRs^24^, including TCRs reactive to *Candida tropicalis* and *Saccharomyces cerevisiae*, were linked to the TCRs identified in our convergent cluster analyses (TCRα: 1.1%, TCRβ: 0.2%). Only 0.1% of the CD-related enhanced TCRβ sequences as reported in the patent^25^ were also identified in our convergent cluster analyses. The eight selected CD-related clusters from the present study almost have no overlap with published TCRs.

In conclusion, a substantial proportion of potentially CD-related TCR sequences from literature were present in the TCR repertoires in the present study but not specifically found to be CD-enriched. The TCR-sequences we identified through cluster analyses were mostly new, i.e. not overlapping with the earlier reported TCRs.

## DISCUSSION

Within the setting of monozygotic twin pairs concordant and discordant for CD, the richness, diversity and clonal expansion of the TCRα and TCRβ repertoire of CD4+ memory gut-homing, non-gut-homing and Tregs was comparable between CD patients, healthy cotwins and healthy controls. Across CD4+memory gut-homing, CD4+ memory non-gut-homing T-cells, and Tregs, the TCR repertoire of Tregs was less diverse and more clonally expanded. Interestingly, we found an increase in clonotype overlap in the TCR repertoire of twin pairs concordant for CD but not for discordant twin pairs, which hints at skewing of the TCR repertoire within CD. Next, we identified potential CD-related TCR sequences and TCR clusters via TCR-convergence and -neighbor enrichment analyses. Most of these TCRs do not yet have known antigen specificity. The majority of TCRs within these CD-related clusters have not previously been linked to CD.

A less diverse and more oligoclonal repertoire in blood and mucosa of CD patients has been reported previously.^5–7^ Furthermore, in a mouse model it has been shown that an oligoclonal Treg repertoire is associated with spontaneous development of intestinal inflammation, that could be suppressed by the transfer of Tregs with a diverse repertoire.^26^ In our study, however, the peripheral TCR diversity and clonality per CD4+ celltype was comparable in our study between CD patients, healthy cotwins, and healthy controls.

Increased overlap in the TCR repertoire of monozygotic twin pairs is expected, based on their shared genetic background, including sharing of HLA alleles^27^ and environmental factors. Studies into the TCR repertoire of healthy monozygotic twin pairs showed increased overlap,^28,29^ although the TCR repertoire per cotwin still is person-specific.^30^ In a previous study into the peripheral TCR repertoire among IBD-discordant and concordant twins, with a low number of CD concordant twins, no increase in overlap was found for CD concordant pairs compared to CD discordant pairs.^15^ This might partly be explained by the fact that healthy cotwins from CD discordant pairs are at increased risk of developing CD,^31^ and might therefore already partly display CD-related features. We did find an increased overlap in the TCR repertoire within CD concordant twin pairs compared to random pairs of healthy unrelated individuals, pointing towards antigen-driven TCR-repertoire skewing in CD.

Highly similar TCR sequences may recognize the same epitopes.^16,32^ We therefore did not only focus on exact TCR clonotypes, but performed convergence and enrichment analyses followed by clustering to identify potentially CD-related TCR patterns. Via this approach we were able to identify 8 distinct clusters of TCRs. Interestingly, the TCR sequences in these enriched clusters are independent from reported CD-related TCR sequences^11,15,24,25^, such as the CAITs. The low percentage of overlap with earlier reported sequences has recently also been noted by Pesesky et al when they compared their identified CD-related enhanced TCRβ-sequences^7^ to the CD-related TCRs from literature.^11,24^ This reflects the complexity of identifying CD-related specific TCRs, but also warrants further exploration of our and other identified CD-related TCRs.

Ultimately the goal is to dissect the role of specific antigens and antigen-specific T-cells in CD. Currently it is impossible to reliably predict the cognate antigen for each TCR in-silico.^33^ However, by using the Immunewatch Detect tool^23^ we were able to predict antigen specificities for some of the identified 8 CD-related TCR clusters based on known epitopes, using existing literature and computational models, including microbial and influenza A antigens. Still, for most TCRs no antigen specificity could be confidently predicted. It is important to bear in mind that this tool (and other TCR-antigen prediction tools) are biased towards viral rather than self epitopes, since the majority of TCR data has been generated in relation to viral infections or vaccination studies. This can also be the explanation why we and Rosati et al.^15^ found TCRs directed against Influenza A viruses to be associated with CD. Microbial antigens could be of particular interest in the pursuit of CD-driving antigens, because alterations in the gut microbiome have been implicated in CD, with microbiome alterations not only observed in established disease,^34^ but also in treatment-naïve CD patients,^35^ people at risk of developing IBD,^17^ and within the prediagnostic phase of CD.^36^ Studies into micro- and mycobiota-reactive T-cells in the setting of CD showed that microbiota-reactive CD4+ T-cells in CD are skewed towards a Th17 phenotype,^37,38^ while for mycobiota-reactive CD4+ T-cells a Th1 skewed phenotype was found.^24^ Unfortunately, TCR sequencing data can currently not be directly linked to the microbial compositon.

Establishing such a connection would ultimately provide crucial insights into the immune-microbiome interactions underlying CD.

Our study has several strengths. By studying the TCR repertoire in monozygotic twins we could compare HLA-identical individuals, with shared genetic and (childhood) environmental factors, reducing the influential factors on the TCR repertoire. We both studied the TCRα and TCRβ repertoire in different CD4+ T-cell subsets increasing the resolution compared to for example whole-blood sequencing. It has been shown that TCRs that are not identical, but sufficiently similar can recognize the same antigen.^16,32^ By not only focusing on exact TCRs but including similarity-based convergence and neighbor enrichment analyses, we were able to identify potentially CD-related TCR clusters rather than relying only on exact clonotype matches or enrichment.

Our study also has some limitations. A limited number of participants was included, which is partly compensated for by the twin-design, however a larger number of participants would have added larger statistical power. The sequencing technique used allows for the TCR sequencing of a large number (in our case 100,000) T-cells per sample. However, it is impossible in bulk sequencing to pair TCRα and TCRβ sequences. A paired T-cell receptor would have enabled a more comprehensive and precise study of the complete TCR repertoire. Lastly, we chose to focus on peripheral CD4+ T-cells, so no information on the CD8+ and the mucosal TCR repertoire in CD was generated.

In conclusion, the increased overlap in the TCR repertoire within CD-concordant monozygotic twin pairs, point towards a role for (antigen-driven) skewing of the TCR repertoire in the pathophysiology of CD. We identified novel CD-related TCR clusters with mostly unknown cognate antigens, that are prime targets for further study into the pathogenesis of CD.

### Grant support

Eelco C. Brand is supported by the Alexandre Suerman program for MD and PhD candidates of the University Medical Center Utrecht, the Netherlands. Romi Vandoren is supported by the Interuniversity Special Research Fund (iBOF) [‘Modulating Immunity and the Microbiome for Effective CRC Immunotherapy’ (MIMICRY) Project]. Vincent Van Deuren is supported by the Research Foundation Flanders [FWO: 1SH6624N].

## Disclosures

ECB is co-applicant on an Investigator Initiated research grant of Pfizer. RV nothing to declare.

LL nothing to declare. VVD nothing to declare. NHS nothing to declare.

AP is an employee of AbbVie and may hold stock or stock options in AbbVie.

PM is co-founder, board director, and shareholder of ImmuneWatch, an immunoinformatics company. ImmuneWatch had no role in study design, data collection and analysis, decision to publish, or preparation of the manuscript.

BO has received research support from Pfizer, Galapagos, Abbvie, Takeda, Ferring and consulting and speaker fees from Galapagos, Takeda, Janssen, BMS, Pfizer, Abbvie and Ferring.

FvW is a speaker and/or consultant for Janssen, Johnson & Johnson, and Takeda and has received grants from Regeneron Pharmaceuticals, Leo Pharma, Sanofi, BMS, Galapagos, and Takeda.

### Writing assistance

No writing assistance was obtained for the present study.

## Acknowledgements

The authors thank all participants. The authors thank the core flow-facility of the CTI for their support in FACS-sorting, Noortje van den Dungen for her help with quality control of the isolated RNA, Bart Müskens and Victor Rijnierse for their help with study visits in the TWIN- study, and Sebastiaan Valkiers for his help with the ClusTCR and SNE method. Figure 1 has been created with www.biorender.com.

## Members of the Dutch TWIN-IBD consortium

In alphabetical order of affiliation.

Bas Oldenburg, MD, PhD, Department of Gastroenterology and Hepatology, University Medical Center Utrecht, Utrecht, The Netherlands

Femke van Wijk, PhD, Center for Translational Immunology, University Medical Center Utrecht, Utrecht, The Netherlands

Eelco C. Brand, MD, Department of Gastroenterology and Hepatology & Center for Translational Immunology, University Medical Center Utrecht, Utrecht, The Netherlands Pieter Honkoop, MD, PhD, Department of Gastroenterology and Hepatology, Albert Schweitzer Hospital, Dordrecht, Zwijndrecht, Sliedrecht, The Netherlands

Rutger J. Jacobs, MD, PhD, Department of Gastroenterology and Hepatology, Alrijne Hospital, Leiden, Leiderdorp, Alphen aan den Rijn, The Netherlands

Bart L.M. Müskens, MSc, Department of Gastroenterology and Hepatology, Amsterdam UMC, Amsterdam, The Netherlands.

Cyriel Y. Ponsioen, MD, PhD, Department of Gastroenterology and Hepatology, Amsterdam UMC, Amsterdam, The Netherlands.

Nanne K.H. de Boer, MD, PhD, Department of Gastroenterology and Hepatology, Amsterdam UMC, Amsterdam, The Netherlands.

Yasser A. Alderlieste, MD, Department of Gastroenterology and Hepatology, Beatrix Hospital, Gorinchem, The Netherlands

Margot A. van Herwaarden, MD, PhD, Department of Gastroenterology and Hepatology, Deventer Hospital, Deventer, The Netherlands

Sebastiaan A.C. van Tuyl, MD, PhD, Department of Gastroenterology and Hepatology, Diakonessen Hospital, Utrecht, The Netherlands

Maurice W. Lutgens, MD, PhD, Department of Gastroenterology and Hepatology, Elisabeth- TweeSteden Hospital, Tilburg, The Netherlands A. C. Janneke van der Woude, MD, PhD, Department of Gastroenterology and Hepatology, Erasmus Medical Center, Rotterdam, The Netherlands

Wout G.M. Mares, MD, Department of Gastroenterology and Hepatology, Gelderse Vallei Hospital, Ede, The Netherlands

Daan B. de Koning, MD, Department of Gastroenterology and Hepatology, Gelre Hospitals, Apeldoorn, Zutphen, The Netherlands

Joukje H. Bosman, MD, Department of Gastroenterology and Hepatology, Groene Hart Hospital, Gouda, The Netherlands

Juda Vecht, MD, PhD, Department of Gastroenterology and Hepatology, Isala, Zwolle, The Netherlands

Anneke M.P. de Schryver, MD, PhD, Department of Gastroenterology and Hepatology, Jeroen Bosch Hospital, Den Bosch, The Netherlands

Andrea E. van der Meulen-de Jong, MD, PhD, Department of Gastroenterology and Hepatology, Leiden University Medical Center, Leiden, The Netherlands

Marieke J. Pierik, MD, PhD, Department of Gastroenterology and Hepatology, Maastricht University Medical Center, Maastricht, The Netherlands

Paul J. Boekema, MD, PhD, Department of Gastroenterology and Hepatology, Maxima Medical Center, Veldhoven, Eindhoven, The Netherlands

Robert J. Verburg, MD, PhD, Department of Gastroenterology and Hepatology, Medical Center Haaglanden, Den Haag, The Netherlands

Bindia Jharap, MD, PhD, Department of Gastroenterology and Hepatology, Meander Medical Center, Amersfoort, The Netherlands

Gonneke Willemsen, PhD, Faculty of Health, Sports and Social Work, Inholland University of Applied Sciences, Haarlem, the Netherlands,

Dorret I. Boomsma, PhD, Department of Complex Trait Genetics, Center for Neurogenomics and Cognitive Research Vrije Universiteit Amsterdam, The Netherlands

Jeroen M. Jansen, MD, Department of Gastroenterology and Hepatology, OLVG Oost, Amsterdam, The Netherlands

Pieter C.F. Stokkers, MD, PhD, Department of Gastroenterology and Hepatology, OLVG West, Amsterdam, The Netherlands

Frank Hoentjen, MD, PhD, Division of Gastroenterology, Department of Medicine, University of Alberta, Edmonton, Canada. Previously: department of Gastroenterology and Hepatology, Radboud University Medical Center, Nijmegen, The Netherlands

Rutger Quispel, MD, Department of Gastroenterology and Hepatology, Reinier de Graaf Gasthuis, Delft, The Netherlands

Carmen S. Horjus Talabur Horje, MD, PhD, Department of Gastroenterology and Hepatology, Rijnstate Hospital, Arnhem, The Netherlands

Paul C. van de Meeberg, MD, PhD, Department of Gastroenterology and Hepatology, Slingeland Hospital, Doetinchem, The Netherlands

Nofel Mahmmod, MD, Department of Gastroenterology and Hepatology, St. Antonius Hospital, Nieuwegein, Utrecht, The Netherlands

Meike M.C. Hirdes, MD, PhD, Department of Gastroenterology and Hepatology, St. Franciscus Gasthuis, Rotterdam, The Netherlands

Rachel L. West, MD, PhD, Department of Gastroenterology and Hepatology, St. Franciscus Gasthuis, Rotterdam, The Netherlands

Marleen Willems, MD, Department of Gastroenterology and Hepatology, St. Jansdal Hospital, Harderwijk, The Netherlands

Itta M. Minderhoud, MD, PhD, Department of Gastroenterology and Hepatology, Tergooi Hosptial, Blaricum, Hilversum, The Netherlands

Herma H. Fidder, MD, PhD, Department of Gastroenterology and Hepatology, University Medical Center Utrecht, Utrecht, The Netherlands

Fiona D.M. van Schaik, MD, PhD, Department of Gastroenterology and Hepatology, University Medical Center Utrecht, Utrecht, The Netherlands

Nynke A. Boontje, MSc, Department of Gastroenterology and Hepatology, University Medical Center Utrecht, Utrecht, The Netherlands

Sarah L. Dijkstra, MD, Department of Gastroenterology and Hepatology, University Medical Center Utrecht, Utrecht, The Netherlands

A. Sophie Bakker, MSc, Department of Gastroenterology and Hepatology & Center for Translational Immunology, University Medical Center Utrecht, Utrecht, The Netherlands Rinse K. Weersma, MD, PhD, Department of Gastroenterology and Hepatology, University Medical Center Groningen, Groningen, The Netherlands

Marielle J.L. Romberg-Camps, MD, PhD, Department of Gastroenterology and Hepatology, Zuyderland Hospital, Sittard-Geleen, Heerlen, The Netherlands

