## Supplementary tables 1-2 and supplementary figure legends for "Clonal overlap and convergent clustering of T-cell receptor signatures in Crohn’s disease in monozygotic twins"

| Supplementary table 1. Antibodies used for flow-assisted cell-sorting and flow cytometry | | | | | | |
| --- | --- | --- | --- | --- | --- | --- |
| Target | **Fluorochrome** | **Channel** | **Clone** | **Dilution** | **Company** | **Cat number** |
| CD45RA | FITC | Blue1 | HI100 | 50 | BD | 555488 |
| CD8a | PerCP-Cy5.5 | Blue3 | RPA-T8 | 1000 | Biolegend | 301032 |
| CCR9 | APC | Red1 | L053E8 | 10 | Biolegend | 358908 |
| CD3 | AF700 | Red 1/1 | UCHT1 | 50 | Biolegend | 300424 |
| Integrin β7 | BV421 | Violet1 | FIB504 | 200 | BD | 564283 |
| Viability dye | eF506 | Violet2 |  | 1000 | eBioscience | 65-2860-40 |
| CD127 | BV605 | Violet3 | A019D5 | 50 | Sony Biotechnology | 2356670 |
| CD25 | BV711 | Violet5 | 2A3 | 50 | BD | 563159 |
| CD4 | BV785 | Violet6 | OKT4 | 100 | Biolegend | 317442 |
| CCR7 | PE | YeGr1 | 3D12 | 25 | eBioscience | 12-1979-42 |
| Integrin α4 | Pe-Cy7 | YeGr5 | 9F10 | 50 | Biolegend | 304314 |
| Only used for FOXP3 check^a^ | | | | | | |
| FOXP3 | eF450 | Violet1 | PCH101 | 50 | eBioschience | 48-4776-42 |
| AF, alexa fluor; APC, allophycocyanin; BV, brilliant violet; eF, eFluor; FITC, fluorescein isothiocyanate; PE, phycoerythrin; Pe-Cy, phycoerythrin-cyanin; PerCP-Cy, peridinin chlorophyll-cyanin.  ^a^The cells used for the FOXP3 check were not stained with Integrin β7, because of the overlap in channel needed for flow cytometry measurement. | | | | | | |

| Supplementary table 2. Baseline and sample characteristics of included patients for flow-cytometry analyses. | | | |
| --- | --- | --- | --- |
|  | Crohn’s disease concordant twin pairs  (6 pairs, 12 individuals) | Crohn’s disease  discordant twin pairs  (4 pairs, 8 individuals) | Healthy controls  (6 individuals) |
| *Demographics & clinical characteristics* | | | |
| Female sex | 4 (33.3%) | 4 (50%) | 3 (50%) |
| Age | 27.5 (23, 55) | 35 (30, 41.8) | 34.5 (35.5, 51.3) |
| Body mass index | 22.3 (21.3, 22.9) | 23.1 (21, 25.3) | NA^a^ |
| Current smoking | 1 (8.3%) | 2 (25%) | NA^a^ |
| Disease phenotype |  |  |  |
| *Crohn’s disease* | 12 (100%) | 4 (50%) | 0 |
| *No inflammatory bowel disease* | 0 | 4 (50%) | 6 (100%) |
| Zygosity |  |  |  |
| *Monozygotic* | 12 (100%) | 8 (100%) | - |
| *Dizygotic* | 0 | 0 | - |
| *Crohn’s disease characteristics^b^* | | | |
| Disease duration (months) | 142 (13, 253) | 104 (17.4, 189) | - |
| Age of diagnosis (Montreal classification) |  |  |  |
| *A1 (≤16 years)* | 2 (16.7%) | 0 | - |
| *A2 (17-39 years)* | 10 (83.3%) | 3 (75%) | - |
| *A3 (≥40 years)* | 0 | 1 (25%) | - |
| Location (Montreal classification) |  |  |  |
| *L1 (ileum only)* | 6 (50%) | 2 (50%) | - |
| *L2 (colon only)* | 3 (25%) | 1 (25%) | - |
| *L3 (ileocolonic)* | 3 (25%) | 1 (25%) | - |
| *L4 (proximal of ileum)* | 2 (16.7%) | 0 | - |
| Behavior (Montreal classification) |  |  |  |
| *B1 (non-stricturing, non-penetrating)* | 5 (41.7%) | 3 (75%) | - |
| *B2 (stricturing)* | 5 (41.7%) | 1 (25%) | - |
| *B3 (penetrating)* | 2 (16.7%)^c^ | 0 | - |
| *p (perianal modifier)* | 0 | 0 | - |
| Harvey-Bradshaw Index^d^ | 2.5 (1.8, 3) | 3 (1, 5) | - |
| Signs of disease activity^e^ | 4 (33.3%) | 2 (50%) | - |
| Current Crohn’s disease medication |  |  |  |
| *No Crohn’s disease medication* | 1 (8.3%) | 1 (25%) | - |
| *5-aminosalicylic acid (5-ASA)* | 0 | 1 (25%) | - |
| *Corticosteroids* | 0 | 0 | - |
| *Methotrexate* | 0 | 0 | - |
| *Thiopurine* | 9 (75%) | 3 (75%) | - |
| *Anti-tumor necrosis factor-α* | 6 (50%) | 0 | - |
| *Vedolizumab (anti-integrin α4β7)* | 0 | 0 | - |
| *Ustekinumab (anti-interleukin12/23)* | 0 | 0 | - |
| NA, not applicable.  Continuous variables are depicted as median (p25, p75), and categorical variables as number (proportion) unless indicated otherwise.  RNA-isolation of 12 samples of 4 Crohn’s disease patients and 2 healthy controls did not pass our preset quality criteria and were therefore not used for T-cell receptor sequencing. Because of a low yield of sorted regulatory T-cells (<30 000) the samples of one Crohn’s disease concordant twin pair was not included for T-cell receptor sequencing. T-cells from 4 extra Crohn’s disease patients and 2 healthy controls were therefore sorted and used for T-cell receptor sequencing. Nonetheless, the flow-cytometry data of these patients was included in our flow-cytometry analyses.  ^a^Data not collected for healthy controls.  ^b^Crohn’s disease characteristics are only relevant to the Crohn’s disease patients and therefore proportions reflect the percentage relative to the total number of Crohn’s disease patients per group.  ^c^One of these patients is classified as B2 and B3.  ^d^The patient Harvey-Bradshaw index is a symptom based score with higher scores indicating more active disease.^1^  ^e^Disease activity was defined as Harvey Bradshaw Index >4 or signs of inflammation as noted during rectoscopy. | | | |

**Supplementary tables 3 & 4**

These tables are displayed in supplementary excel files.

**SUPPLEMENTARY FIGURE LEGENDS**

**Supplementary figure 1. Gating strategy and FOXP3 check.** A) A representative example of the gating strategy for PBMCs is shown. CD4+ Tregs (alive CD3+CD8-CD4+CD25+CD127low), gut-homing memory CD4+ T-cells (alive CD3+CD8-CD4+CD25-CCR7+/-CD45RA-integrinα4β7+), and non-gut-homing memory CD4+ T-cells (alive CD3+CD8-CD4+CD25-CCR7+/-CD45RA-integrinα4β7-) were FACS-sorted. These T-cell subsets were subsequently used for TCRα and TCRβ repertoire analyses. B) A subsample of sorted Tregs and non Tregs (alive CD3+CD8-CD4+CD25-) were intracellularly stained for transcription factor FOXP3, to check the efficacy of the Treg sort. Here, we show a representative histogram of the fluorescence for the FOXP3 directed antibody on FACS, showing a distinct difference in FOXP3 expression between non-Tregs (no expression) and Tregs (high expression) as expected. The FOXP3 check confirmed the ability of our sorting strategy to separate Tregs (median proportion FOXP3 positive: 87.9% measured in 16 samples) from non-Tregs (median proportion FOXP3 negative: 98.5%, measured in 16 samples).

AF, alexa fluor; BV, brilliant violet; eF, eFluor; FACS, fluorescence-activated cell sorting; FITC, fluorescein isothiocyanate; FOXP3, forkhead box P3; PBMCs, peripheral blood mononuclear cells; PE, phycoerythrin; Pe-Cy, phycoerythrin-cyanin; PerCP-Cy, peridinin chlorophyll-cyanin; Tregs, regulatory T-cells.

**Supplementary figure 2. Study overview of included TCR-samples**.

The unique number of T-cell receptor sequences found per cell-type and per sample are displayed, as well as the proportion of the total number of T-cell receptor sequences (i.e. unique T-cells) per cell type.

CC, concordant Crohn’s disease; DC, discordant Crohn’s disease; gut-homing CD4+ memory T-cells, alive CD3+CD8-CD4+CD25-CCR7+/-CD45RA-Integrinα4β7+ T-cells; non gut-homing CD4+ memory T-cells, alive CD3+CD8-CD4+CD25-CCR7+/-CD45RA-Integrin-α4β7- T-cells; TCR, T-cell receptor; Tregs, regulatory T-cells (alive CD3+CD4+CD8-CD25+CD127low T-cells)

**Supplementary figure 3. Peripheral CD4+ T-cell composition.** A) Stacked bar chart showing the composition of peripheral non-Tregs. No statistically significant differences were found between Crohn’s disease twins, healthy cotwins and healthy controls in the proportions of memory (alive CD3+CD4+CD8-CD25-CD45RA-) and naive (alive CD3+CD4+CD8-CD25-CD45RA+CCR7+) CD4+ T-cells. Distribution of B) regulatory T-cells (alive CD3+CD4+CD8-CD25+CD127low), C) gut-homing CD4+ memory T-cells (alive CD3+CD8-CD4+CD25-CCR7+/-CD45RA-Integrinα4β7+), and D) non gut-homing CD4+ memory T-cells (alive CD3+CD8-CD4+CD25-CCR7+/-CD45RA-Integrin-α4β7-) between Crohn’s disease twins, healthy cotwins and healthy controls.

MZ, monozygotic; N, number of participants, TCR, T-cell receptor; temra, terminally differentiated effector memory T cells; Tregs, regulatory T-cells.

**Supplementary figure 4. TCR convergence colored per cell type.** Same figures as shown in figure 4B and 4C, but now colored per cell subtype.

Gut-homing CD4+ memory T-cells, alive CD3+CD8-CD4+CD25-CCR7+/-CD45RA-Integrinα4β7+ T-cells; non gut-homing CD4+ memory T-cells, alive CD3+CD8-CD4+CD25-CCR7+/-CD45RA-Integrin-α4β7- T-cells; TCR, T-cell receptor; regulatory T-cells, alive CD3+CD4+CD8-CD25+CD127low T-cells.

**Supplementary figure 5.** **TCR neighbor enrichment analyses and clustering more detailed.** TCR neighbor enrichment is calculated comparing the number of neighbor TCR sequences (i.e. similar TCR sequences) within one TCR repertoire compared to a synthetic background repertoire. Only statistically significantly (Bonferroni adjusted P-value ≤ 0.05) enriched TCRs are displayed separately for Crohn’s disease patients, healthy cotwins and healthy controls. Results are colored by A) TCRα and TCRβ, B) cell type, C) predicted epitope based on Immunewatch Detect^2^, D) the individual in which the TCRs are identified in. Within the Crohn’s disease patients a few TCRs of non-gut homing CD4+ memory T-cells which are partly predicted to be directed against viral and partly against microbial antigens are the most statistically significantly enriched TCRs. The TCRs with the lowest p-value all come from one Crohn’s disease patient. Panel A-D show the same analyses as figure 5A in which the results are presented in a more condensed form.

CC, concordant Crohn’s disease; DC, discordant Crohn’s disease; Gut-homing CD4+ memory T-cells, alive CD3+CD8-CD4+CD25-CCR7+/-CD45RA-Integrinα4β7+ T-cells; Log2FC, log2fold change; non gut-homing CD4+ memory T-cells, alive CD3+CD8-CD4+CD25-CCR7+/-CD45RA-Integrin-α4β7- T-cells; TCR, T-cell receptor; regulatory T-cells, alive CD3+CD4+CD8-CD25+CD127low T-cells.

**Supplementary figure 6.** **Example of a potentially CD-related TCR cluster**. These clusters are identified through TCR convergence and patient-level neighbor enrichment analyses of which 8 clusters are identified as the potentially most relevant CD-related clusters (table 2 and supplementary table 3). A) plot that shows the cluster analyses as also shown in figure 4D highlighting cluster 1 [CASS.[GT]gnYGYTF] B) The number of TCRs within this cluster as found per participant in the study. C) The most shared TCRs among participants within this cluster. D) Bar graph showing number of TCRs within this cluster per T-cell subtype (i.e. CD4+ gut-homing memory T-cells, CD4+ non gut-homing memory T-cells, and CD4+ Tregs). E) Boxplot showing the absolute clonecount per TCR from this cluster separated for CD patients, healthy cotwins and healthy controls. For none of the TCRs of this cluster a confident epitope prediction could be made by Immunewatch Detect^2^.

CD, Crohn’s disease; Gut-homing CD4+ memory T-cells, alive CD3+CD8-CD4+CD25-CCR7+/-CD45RA-Integrinα4β7+ T-cells; non gut-homing CD4+ memory T-cells, alive CD3+CD8-CD4+CD25-CCR7+/-CD45RA-Integrin-α4β7- T-cells; CD4+ regulatory T-cells, alive CD3+CD4+CD8-CD25+CD127low T-cells; TCR, T-cell receptor; Tregs, regulatory T-cells.
