## Supplementary figures and images for "Clonal overlap and convergent clustering of T-cell receptor signatures in Crohn’s disease in monozygotic twins"

### Supplementary figure 1

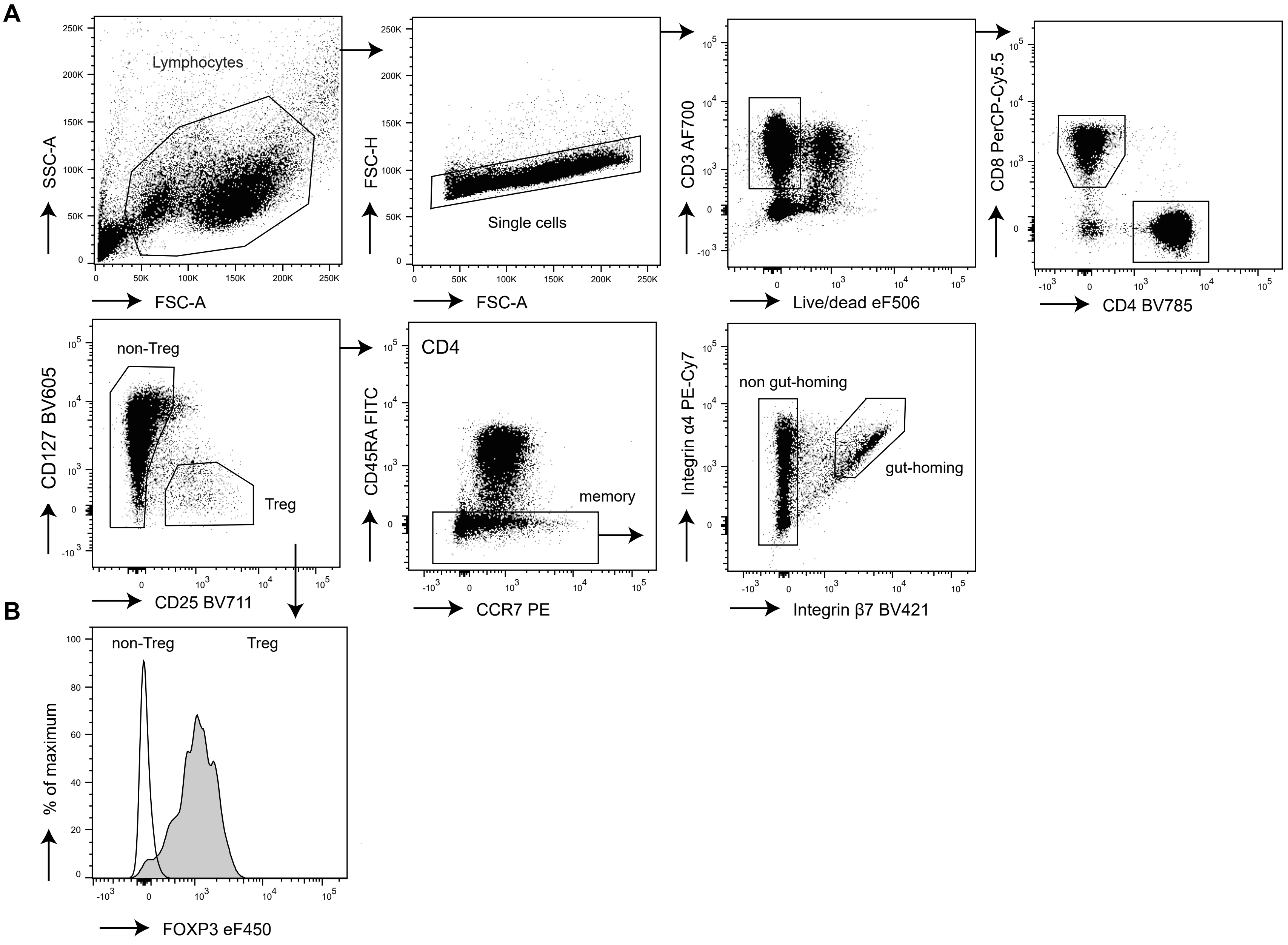

### Supplementary figure 2

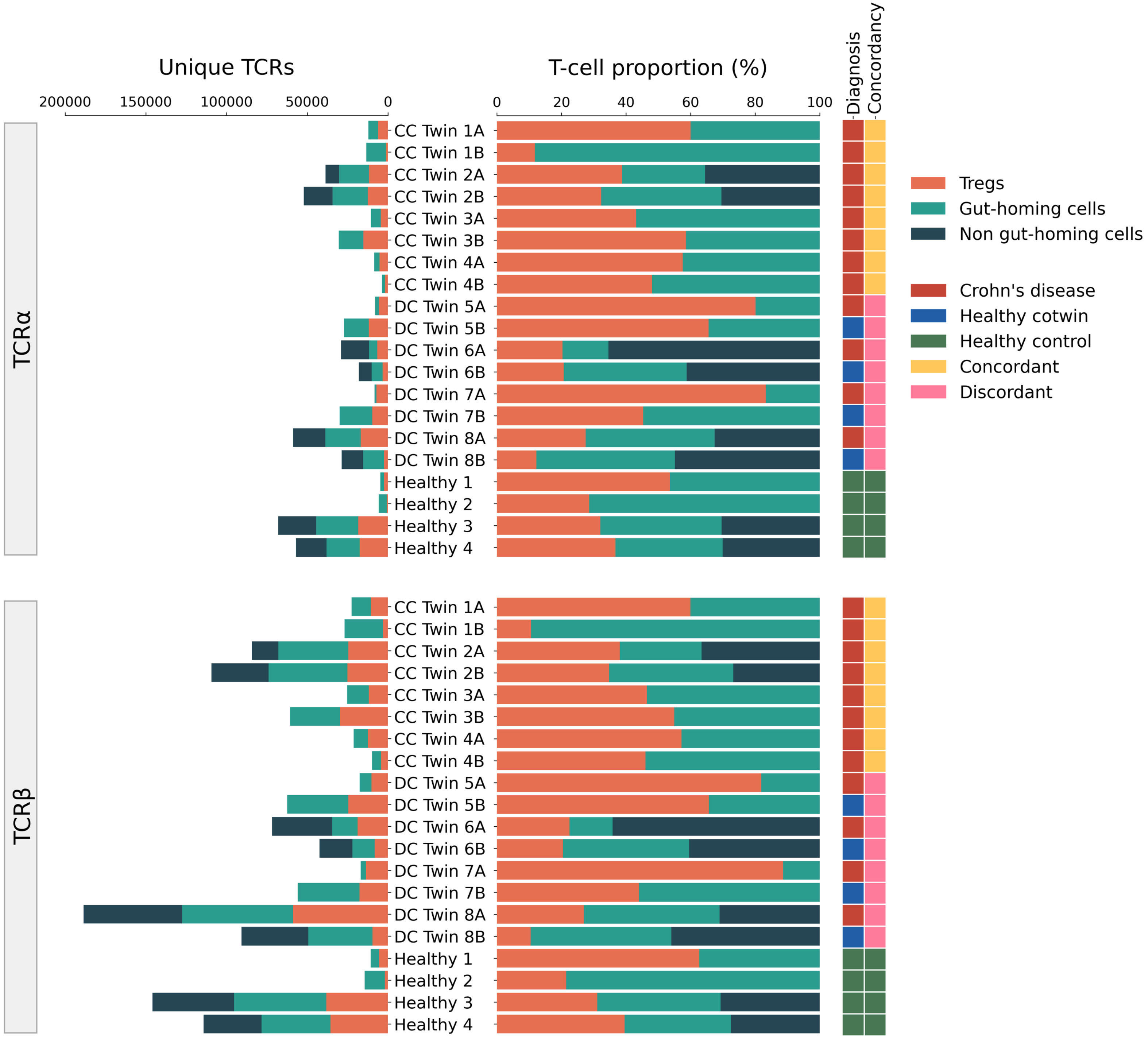

### Supplementary figure 3

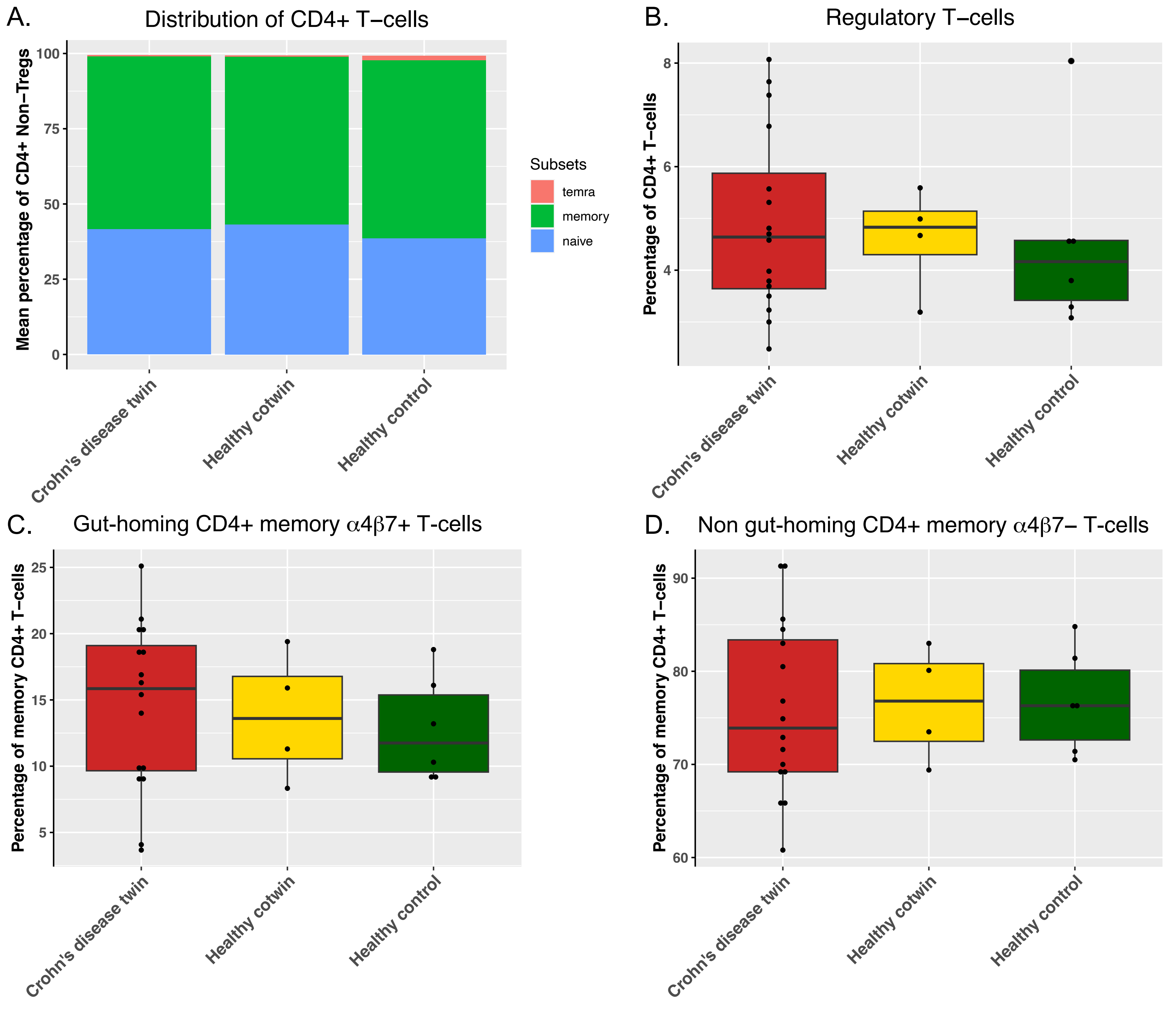

### Supplementary figure 4

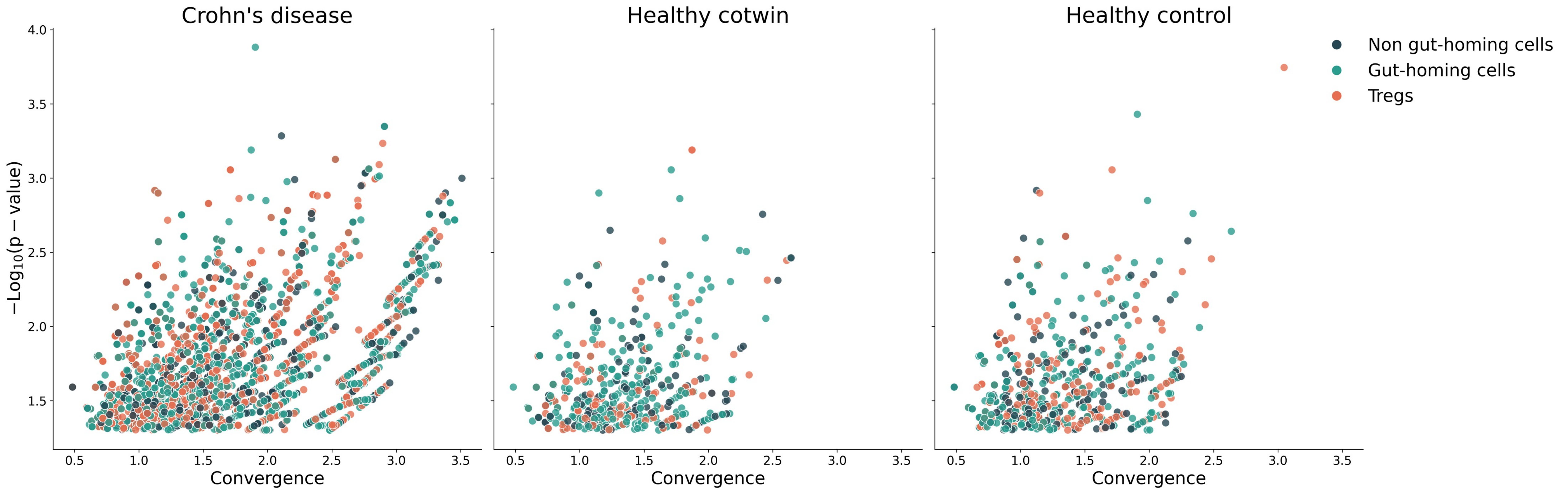

### Supplementary figure 5

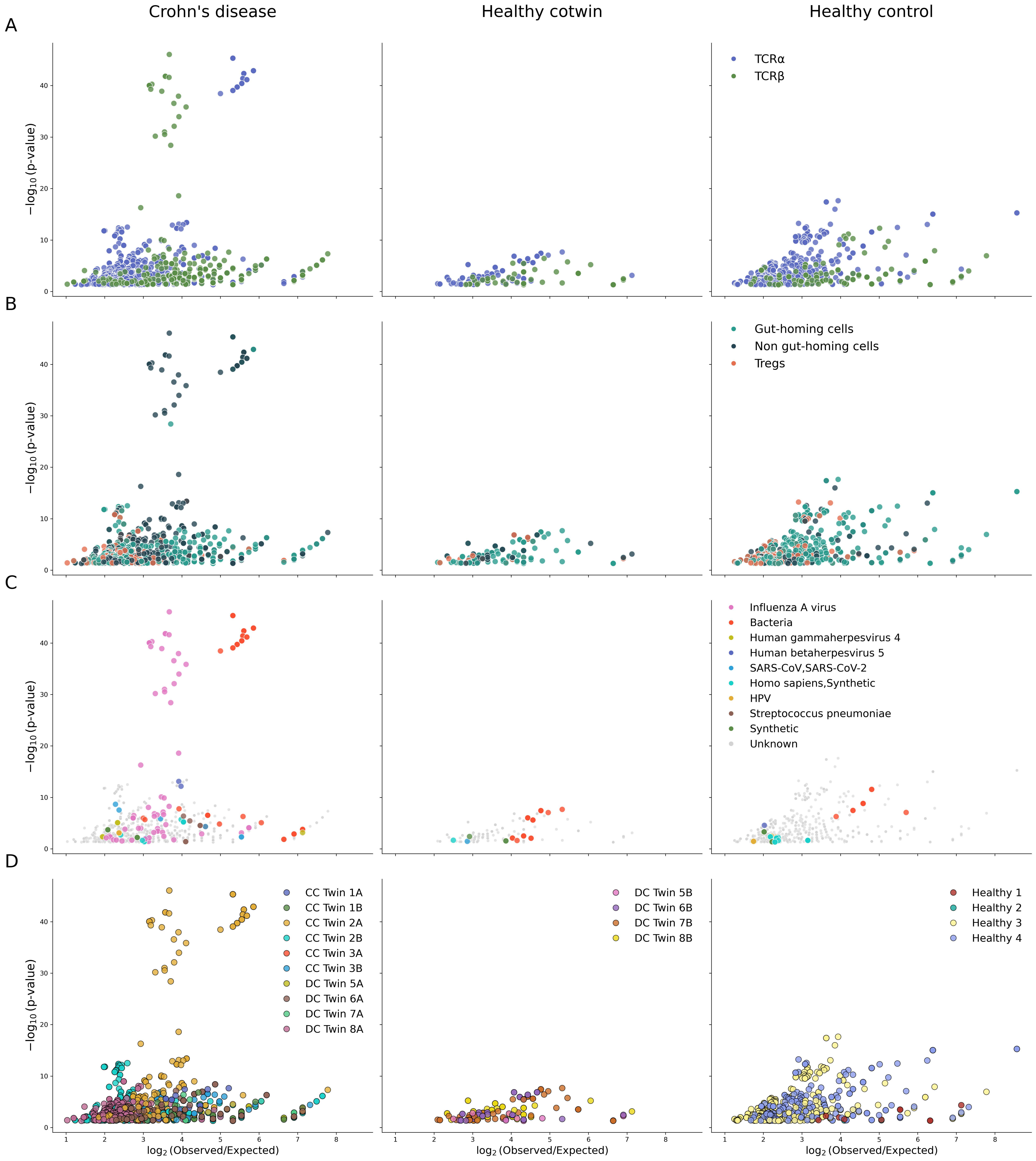

### Supplementary figure 6

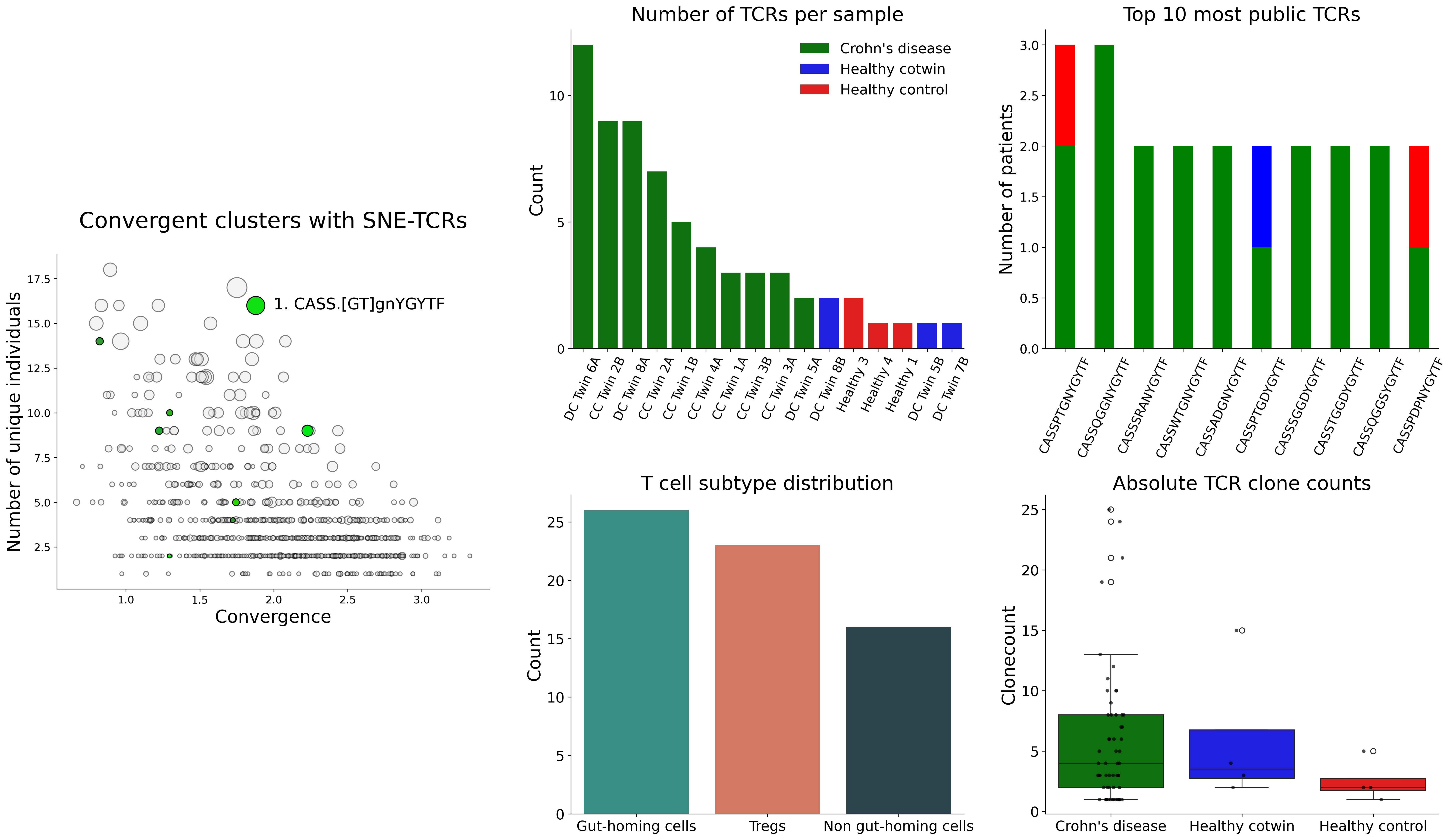
